# Intrinsic spinal cord circuits compare sensory inputs and efference copies to correct for perturbations to ongoing movement

**DOI:** 10.1101/2025.08.27.672590

**Authors:** M. Gorkem Ozyurt, Sarah Chiasson, Alex M Laliberte, Filipe Nascimento, Emam Khan, William P. Mayer, Gardave S. Bhumbra, Turgay Akay, Tuan V. Bui, Marco Beato, Robert M. Brownstone, Rémi Ronzano

**Affiliations:** Department of Neuromuscular Diseases, University College London, London, United Kingdom; Department of Neuroscience Physiology and Pharmacology (NPP), University College London, Gower Street, WC1E 6BT London, UK; Brain and Mind Research Institute, Department of Biology, University of Ottawa, Ottawa, ON, Canada; Atlantic Mobility Action Project, Brain Repair Centre, Department of Medical Neuroscience, Dalhousie University, Halifax, Nova Scotia, Canada; Department of Medical Neuroscience, Dalhousie University, Saint John, New Brunswick, Canada

## Abstract

Successful movement requires continuous adjustments in response to changes in internal and external environments. To do so, neural circuits continuously compare efference copies of motor commands with sensory feedback to respond to sensory prediction errors. Some responses need to be very fast, indicating that they likely occur within spinal cord circuits. Here, we describe spinal cord circuits involving dI3 neurons, showing that they receive multimodal sensory inputs and direct efferent copies from both Renshaw cells and motor neurons. We further show that they form connections to motor pools. Reducing dI3 neuronal activity diminished stumbling responses, as did disrupting Renshaw cell circuits, providing evidence for a comparator role of dI3 neurons for online corrections. Together, our findings reveal a pivotal role for dI3 neurons functioning as comparators of efference copies and external sensory feedback to mediate rapid corrections of ongoing movements.

## Introduction

Nervous systems evolved to take corrective actions in response to unexpected perturbations during ongoing movement. Moreover, to avoid potentially devastating consequences, corrections are often needed with minimal delay. For example, without immediate corrective action, an unexpected obstacle during gait could lead to a fall and result in the loss of prey, becoming prey, or as is the case for humans, serious injury or death. In fact, falls are the second leading cause of accidental death in the world, with 79 people dying *per hour* from falls, and many others (about 4,250 per hour) being significantly injured^1^. It is thus crucial to understand how nervous system circuits are wired in such a way that the incidence of serious errors is minimised.

Principles of movement correction have previously been outlined, with control theory providing a solid foundation^2^. These principles can be applied over many time scales – from milliseconds to days. The fundamental mechanisms underlying motor adaptation and learning have been extensively studied in cerebellar systems^3,4^. The principle outlined is that neurons called “comparators” are responsive to unexpected sensory inputs. To do so, these neurons need to “know” the sensory consequences of the intended movement (reafferent signals) and calculate sensory prediction errors to optimize the completion of the movement^5–9^. These predicted sensory consequences are derived from the motor program. That is, comparator neurons continually use an inhibitory model derived from an efference copy of the motor command and compare this input (which “predicts” reafferent input) with the actual sensory feedback, which includes both reafferent and exafferent (not resulting from the motor program) signals (“instructive” input).

While the brain is undoubtedly key for adaptation and learning, it may be too remote (in distance and synapses) from the centres that control spinal motor neurons to immediately (in milliseconds) correct fundamental errors during ongoing movement^10^. On the other hand, spinal cord circuits have the capacity to integrate multimodal sensory feedback to generate immediate motor responses through corrective reflexes^11^. But for neurons in these circuits to serve as comparators that respond to unpredicted perturbations, they would also need to receive inputs derived from efference copies. These inputs, particularly during gait – a movement generated by spinal cord circuits – would likely need to originate within the spinal cord for immediate corrections to occur. In many such instances, for example when faced with a stumble, the generated corrective signals depend on the phase of the locomotor cycle and occur with delays of only milliseconds^12–14^. Therefore, there must be spinal cord circuits that integrate information about motor activity – derived from efference copies from local circuits – with sensory feedback to provide movement-appropriate corrections.

We reasoned that comparator neurons in spinal cord circuits would use high fidelity, local information – for example, related to the activity of each muscle – to subtract reafferent signals resulting from activity of local circuits such as those responsible for locomotion. Furthermore, the highest fidelity efference copy would arise from motor neurons, with a neuron to produce a negative “image” intercalated between motor neuron axons and the comparator neurons. Such a spinal circuit was described decades ago, with recurrent branches from motor axons innervating a population of local inhibitory neurons, Renshaw cells^15,16^. We therefore reasoned that Renshaw cells may be responsible for predictive inputs to comparator neurons. But what would be the neural substrate for comparison?

We have previously shown that dI3 neurons – spinal neurons born in the third dorsal spinal progenitor domain and defined by their expression of the transcription factor *Isl1*^17^ – mediate grasp reflexes and are important for plasticity of spinal circuit function following spinal cord transection^18,19^. A subset of this cardinal class receives inputs from locomotor circuits and at least some of them innervate motor neurons^18,20^. Therefore, dI3 neurons are well positioned as candidates to integrate predictive signals derived from efference copies with instructive, sensory input, and thus to provide the substrate for dynamic corrections of ongoing movements. But do they receive a signal predicting reafference, and are they positioned in circuits such that they can function as comparator neurons responsible for error correction?

To map dI3 neuronal circuitry and assess their function in corrective responses, we used a combination of mouse genetics, viral tracing, physiological methods, and behavioural challenges. We show that individual dI3 neurons within an interposed region in the intermediate laminae receive instructive inputs from multimodal sensory modalities as well as inputs from Renshaw cells, and project to flexor, extensor, or both sets of motor neurons. We then show that dI3 neurons mediate corrective responses. Lastly, we provide converging evidence that the direct input from Renshaw cells to dI3 neurons may function to facilitate these immediate corrections of movements. We propose that Renshaw cells thus encode the efference copy transmitted by motor axons to predict a reafferent signal. Taken together, we suggest that a subset of dI3 neurons functions as spinal cord comparator neurons to facilitate fast corrective actions, extending the definition of comparator systems from the brain to local spinal circuits.

## Results

### dI3 neurons receive multimodal sensory feedback segregated along their somato-dendritic compartment

Having previously shown that dI3 neurons receive direct primary afferent input from low threshold cutaneous and proprioceptive sensory neurons^18^, we now sought to further define the distribution of primary afferent inputs to dI3 neurons. In the lumbar cord, dI3 neurons are distributed in the intermediate laminae from the medial to lateral grey matter (Supplemental figure 1A-C, ^18^), with the most medial population restricted to the caudal lumbar cord (Supplemental figure 1D). A large proportion of dI3 neurons are located in a zone previously defined as a region of convergence^21^, through which proprioceptive afferents from multiple muscles project (Figure 1A). We therefore sought to determine whether there was preferential targeting of dI3 neurons from proprioceptive afferents in this region. To do so, we used vGlut1, a marker of low-threshold primary afferents and corticospinal projections, together with parvalbumin (PV) that labels proprioceptive afferents in the postnatal spinal cord^22^.

**Figure 1:**
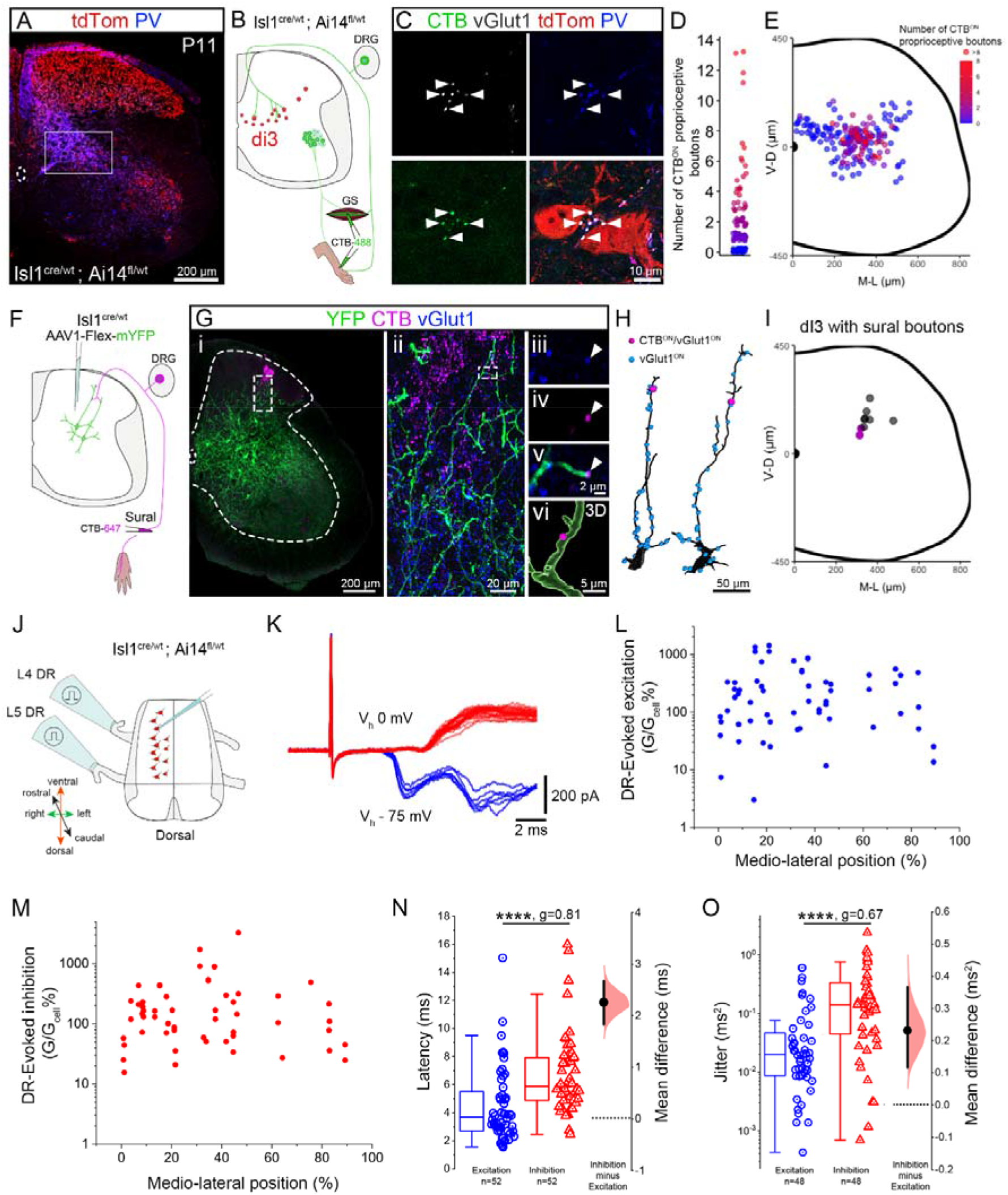
Interposed dI3 neurons receive spatially segregated multimodal sensory afferents and inhibitory inputs from sensory relays. (A) Representative example of a lumbar transverse section of a Isl1^cre/wt^; Ai14^fl/wt^ P11 mouse. The box highlights the region of convergence of proprioceptive afferents where many dI3 neurons are localized. (B) Experimental strategy used to label proprioceptive and cutaneous boutons from the distal hindlimb with CTB injections in the GS and hind paw. (C) Example of two neighbouring dI3 neurons (tdTom^ON^) with proprioceptive boutons labelled from the distal hindlimb (CTB^ON^-vGlut1^ON^-PV^ON^) apposed to their soma. (D) Number of proprioceptive boutons from the distal hindlimb (CTB^ON^-vGlut1^ON^-PV^ON^) apposed to dI3 neurons soma. (E) Spatial distribution of dI3 neurons colour coded by the number of somatic proprioceptive boutons traced from the distal hindlimb. (F) Experimental strategy used to label the dendritic arborization of lumbar dI3 neurons and mechanosensory neurons terminals from the sural nerve. (G) Representative example of a lumbar transverse section from a P35 mouse with terminal endings from the sural nerve labelled with CTB (in magenta), dI3 neurons labelled by membrane tagged YFP (in green) and vglut1 (in blue). The region within the dashed box in (i), is shown at higher magnification in (ii). The boxed region in ii is expanded in iii-v, showing a CTB^ON^-vGlut1^ON^ bouton apposed to a dorsal dendrite of a dI3 neuron and shown in a 3D reconstruction in vi. (H) Examples of reconstructed dI3 dorsal dendrites with apposed vGlut1^ON^ (in blue) and CTB^ON^-vGlut1^ON^ (in magenta) terminals. (I) Position of dI3 neurons with terminals from the sural nerve (CTB^ON^-vGlut1^ON^) apposed to their dorsal dendrites. Dots in magenta correspond to the two examples showed in (H). (J) Schematic of the spinal cord preparation that was used to record dorsal root (DR)-evoked excitation and inhibition (K) of dI3 neurons which were held at -75 mV and 0 mV in voltage-clamp configuration to record excitatory and inhibitory responses respectively. (L-M) Distribution of the size of DR-evoked excitation (L) and inhibition (M) based on the location of dI3 neurons along medio-lateral axis. Inset in (L) describes the coordinate system used to measure the medio-lateral position of recorded neurons. (N-O) Comparisons of inhibition and excitation (N) latency and (O) jitter, where n indicates the number of root stimulations (L4 and L5) plotted. (A-E) Tissues from N=3 animals at P11,n =192 dI3 neurons were analysed, (F-I) Tissues from N=2 animals at P35. (J-N) N=5 mice (P6-11), stimulation of L4 and L5, and recording from n=37 dI3 neurons, plotted values in (N-O) are restricted to root stimulations that triggered both excitatory and inhibitory responses in the recorded dI3 neurons where (N) latency and (O) jitter were measurable, to allow non-hierarchically bootstrapped paired samples Wilcoxon tests and paired Hedge’s G.

We found that more than 80% of dI3 neurons were apposed by a variable number of PV^ON^/vGlut1^ON^ somatic proprioceptive synaptic boutons (Supplemental figure 1E-G). We further observed the greatest density of somatic proprioceptive boutons in the interposed position along the medio-lateral axis, the region of convergence of proprioceptive axons. Since Isl1 is also expressed in the vast majority of sensory neurons, but not in corticospinal neurons^18,23^, we further characterised vGlut1^ON^/tdTom^ON^/PV^OFF^ (in isl1-cre;tdTom mice) boutons as terminals from putative cutaneous sensory neurons. We observed that cutaneous boutons were rarely found on the soma of dI3 neurons, with about 7% of dI3 neurons (13/192) having somatic cutaneous boutons. Furthermore, dI3 neurons showing somatic cutaneous afferents were located in the most medial or dorso-lateral part of the dI3 neuron distribution and largely devoid of somatic proprioceptive boutons (Supplemental figure 1F, H).

To target more specifically sensory neurons from the distal limb, we next traced sensory axons retrogradely using cholera toxin subunit B (CTB). Co-injections were performed into the gastrocnemius muscle and subcutaneously into the hind paw to visualize both proprioceptive and cutaneous fibres (Figure 1B). We found that proprioceptive boutons (CTB^ON^/vGlut1^ON^/PV^ON^) were again mostly restricted to the somata of dI3 neurons in the interposed area (Figure 1C-E).

Conversely, cutaneous afferents labelled from the hind paw were rarely found on dI3 neuron somata with only 2/192 dI3 showing CTB^ON^/vGlut1^ON^/PV^OFF^ boutons, consistent with the position of dI3 neurons being ventral to the region where cutaneous afferents have been shown to terminate^24,25^. Yet we previously showed that dI3 neurons receive afferent input from low-threshold mechanoreceptors and mediate low-threshold cutaneo-motor disynaptic reflexes^18^. We therefore hypothesized that the different modalities of sensory afferents could segregate onto specific subcellular domains of dI3 neurons with the cutaneous afferents projecting specifically to their dorsally-projecting dendrites. To test this hypothesis, we combined intraspinal injection of AAV1-flex-mYFP to induce the expression of a membrane tagged YFP for visualization of dI3 dendritic arbours with retrograde tracing from the sural nerve to label cutaneous afferents (Figure 1F). We first observed that while the vast majority of dI3 somata are restricted to lamina V-VII (Supplemental figure 1A-C, ^18^), their dendrites extend dorsally, densely invading lamina III-IV (with few dendrites even reaching lamina II; Figure 1G). While only 1/50 dI3 somata had cutaneous boutons from the sural nerve in apposition, 100/101 boutons apposed to dI3 neurons were localized on their dorsal dendrites (n=2 animals, Figure 1G-H). Therefore, cutaneous inputs are segregated to the dorsal dendrites of dI3 neurons including dendrites of the interposed dI3 neurons that also receive somatic proprioceptive inputs (Figure 1I).

We then studied functional connectivity of primary afferents to dI3 neurons through patch-clamp recording of dI3 neurons during dorsal root stimulation at low threshold intensities in an isolated spinal cord preparation. Cross-sectional area of dI3 neurons seemed to increase from medial to lateral, but input resistance and whole cell capacitance were not associated with their position (Supplemental figure 1I-K). Following DR stimulation, we observed both excitatory (36/37) and inhibitory (32/37) responses in recorded dI3 neurons. The amplitudes of both conductances (Figure 1J, Supplemental figure 1L-M) were similar to each other and similar along the medio-lateral axis (Figure 1J-L). Both the latency and the jitter of excitation were lower than those of the inhibitory responses recorded (Figure 1M-N), with some of the excitatory responses (even at this young age and room temperature) clearly in the monosynaptic window in keeping with our anatomical observations (Supplemental figure 1N, ^26^).

In summary, proprioceptive sensory inputs densely innervate the somata of dI3 neurons interposed along the medio-lateral axis, whereas cutaneous inputs target the dendrites of these neurons. Furthermore, stimulation of mixed proprioceptive (from flexor and extensor muscles) and low threshold cutaneous afferents leads to both direct excitation and indirect (likely disynaptic) inhibition of these neurons.

### dI3 neurons receive inputs derived from efferent copies of motor outputs

While sensory feedback is essential for motor control, and several spinal populations of dorsal neurons have been shown to rely on mechanosensory afferents to modulate corrective reflexes^27–29^, the integration of processed efferent copies with sensory feedback improves the ability of sensory-motor circuits to estimate the current state of limbs and dynamically adapt the task while avoiding instabilities^3,5,30–32^. To do so, the input derived from efferent copies would be a “negative image” of the efferent command^33,34^. We previously showed that dI3 neurons receive inhibitory inputs from spinal locomotor circuits and suggested that these inputs could represent an efference copy transmitted to dI3 neurons^19^.

We now sought to determine whether dI3 neurons integrate inputs from efference copies with those from sensory afferents and aimed to identify their origin. Efference copies, or copies of the motor program, arise from many levels of the motor circuit hierarchy; the copy with the highest fidelity and most directly related to movement is that carried by motor neuron axons. Therefore, the ideal candidates to convey a negative image would be Renshaw cells (a subpopulation of V1 neurons, which express En1; ^35^), as these inhibitory neurons receive monosynaptic input from motor neurons^36^.

Using En1^cre/wt^ mice crossed with Ai34d^fl/wt^ mice to induce tdTomato expression at the presynaptic terminals of all V1 interneurons (Figure 2A), we could visualize putative Renshaw cell axon boutons by their expression of calbindin and tdTomato (Figure 2B, ^37,38^). We found that about 50% of dI3 neurons had tdTom^ON^/Calbindin^ON^ boutons apposed to their somata, and that these boutons were mostly restricted to the interposed region of the dI3 population (region of convergence) as well as those ventral to this region (Figure 2C-D). Since Renshaw cells are not the only V1 interneurons expressing calbindin in the spinal cord^39^, and anatomically they can be identified solely by using the intersection between their expression of calbindin and their position at the ventral edge of the grey matter, we performed monosynaptic retrograde rabies virus tracing from dI3 neurons using helper constructs with a nuclear GFP reporter for the glycoprotein G and a nuclear BFP reporter to identify cells positive for the avian TVA receptor followed 2 weeks later by an intraspinal injection of RABV-N2c-EnvA-ΔG-mCherry (Figure 2E). We first confirmed that only dI3 neurons and none of the motor neurons or proprioceptive (PV^ON^) sensory neurons (which also express Isl1) were starter cells (BFP^ON^GFP^ON^mChe^ON^, Figure 2F, Supplementary figure 2A-B). Then, using the spatial location of traced neurons combined with calbindin expression to identify Renshaw cells, we observed mChe^ON^ retrogradely-labelled Renshaw cells in 3 out of 4 samples demonstrating that indeed Renshaw cells provide direct inputs to dI3 neurons (Figure 2F-G).

**Figure 2:**
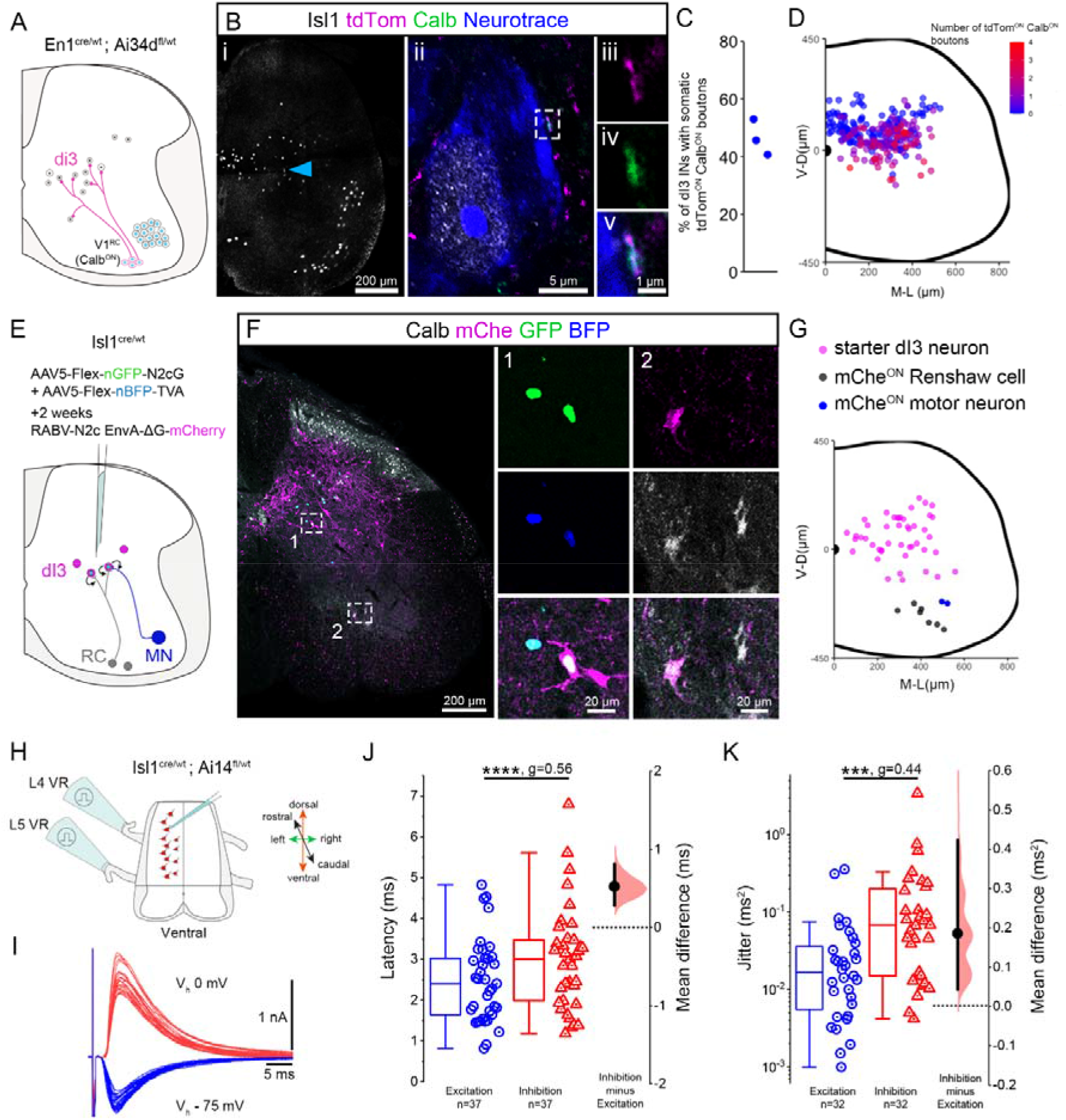
dI3 neurons receive efferent copies from Renshaw cells and motor neurons. (A) Experimental strategy to visualize presynaptic boutons from Calbindin^ON^ V1 interneurons on dI3 somata. (B) Representative example of a lumbar transverse section showing Isl1 (in grey) at P8. The blue arrowhead highlights the dI3 neuron showed in (ii) at a higher magnification with a Calbindin (in green), tdTomato (in magenta) double positive boutons apposed to its soma (neurotrace staining in blue). The dashed box highlights the presynaptic boutons at higher magnification in (iii-v) with the colocalization of calbindin and tdTomato apposed to the membrane (neurotrace staining) of a dI3 neuron. (C) Percentage of dI3 neurons (Isl1^ON^) with Calbindin^ON^ V1 boutons apposed to their soma. Each data point represents the average of one animal. (D) Spatial distribution of dI3 neurons colour coded by the number of Calbindin^ON^ V1 boutons apposed to their soma. (E) Schematic of the method used to trace presynaptic partners to dI3 neurons. The two helper AAVs were injected in lumbar cord of juvenile Isl1^cre/wt^ mouse followed two weeks later by the injection of CSV-N2c-EnvA-ΔG-mChe. (F) Example of a lumbar transverse hemisection showing a mChe^ON^GFP^ON^BFP^ON^ starter dI3 neuron (dashed box 1, at higher magnification on the right side) and a mChe^ON^Calb^ON^ Renshaw cell (dashed box 2, at higher magnification on the right side). (G) Spatial distribution of starter dI3 neurons (in magenta), mChe^ON^Calb^ON^ Renshaw cell presynaptic to dI3 neuron (in grey) and mChe^ON^ChAT^ON^ motor neuron presynaptic to dI3 neuron (in blue). (H) Schematic of the longitudinal preparation of the ventral lumbar cord used to record recurrent excitation and inhibition from dI3 neurons. (I) Sample traces of voltage clamp recording showing recurrent inhibition (red, holding potential 0 mV) and excitation (blue, holding potential -75 mV). (J-K) Comparisons of inhibition and excitation (J) latency and (K) jitter following VR-stimulation where n indicates the number of L4 and L5 responses plotted. (C-D) N=3 animals between P7-P8, n= 273 dI3 neurons analysed. (G) Overlap of spatial mapping from N=4, 6 weeks old animals in which starter cells were mapped on all slices while Renshaw cells and motor neurons were mapped on every other slices. (J-K) N=8 mice (P5-15), L4 and L5 roots stimulated and n=40 dI3 neurons, plotted values are restricted to root stimulations that triggered both excitatory and inhibitory responses in the recorded dI3 neurons where (J) latency and (K) jitter were measurable to allow hierarchically (J) and non-hierarchically (K) bootstrapped paired samples Wilcoxon tests and paired Hedge’s G analysis.

In the samples with the most efficient tracing, we also observed 2 labelled motor neurons showing that dI3 neurons also receive direct collaterals from motor neurons (Figure 2G, Supplementary figure 2C). The low number of motor neurons labelled suggest that these connections may be more sparse than those from Renshaw cells.

Our anatomical observations showed the existence of direct monosynaptic connections from Renshaw cells and, to a lesser extent, from motor neurons to dI3 neurons. However, by anatomy alone, we could not assess the density and strength of these connections. We therefore proceeded to record dI3 neurons identified by tdTomato expression while stimulating ventral roots (VR) to activate motor axons (Figure 2H). In our reduced (isolated ventral lumbar spinal cord) preparations, we found that about 75% (n=30/40) of dI3 neurons received VR-evoked inhibition of varying magnitude (Figure 2I, Supplemental figure 2E-F). Further, these responses showed a low latency and jitter consistent with disynaptic connectivity (Figure 2J-K).

In addition, the majority of dI3 neurons (88%, n=35/40) also received VR-evoked excitation (purely glutamatergic) that was, for a given dI3 neuron, of consistently lower amplitude than inhibition (Figure 2I, Supplemental figure 2D-F, J-K). The monosynaptic nature of these excitatory inputs was supported by their capacity to follow high frequency stimulation, their shorter latency, and lower jitter relative to that observed for inhibition, with most of the excitatory responses falling into or close to the clear monosynaptic window^26^ especially when taking into account the effect of age on these parameters (Figure 2J-K; Supplemental figure 2G-I; ^40^).

To corroborate whether the VR-evoked inhibition of dI3 neurons was indeed mediated by Renshaw cells, a non-competitive antagonist of nicotinic acetylcholine receptors, mecamylamine, was used to block nicotinic transmission from motor neurons to Renshaw cells,^41,42^ leaving intact the purely glutamatergic inputs from motor neurons to dI3 neurons (Supplementary Figure 2J-K). After mecamylamine application, Renshaw cell spiking activity (cell-attached recordings) following VR-stimulation was either abolished (6/10 cells) or the latency and jitter increased significantly (4/10 cells, Supplementary figure 2L-N). We found that mecamylamine led to a 35% decrease of the inhibitory currents recorded from dI3 neurons following VR stimulation, similar to the decrease of recurrent inhibition of motor neurons in response to VR stimulation (∼25%, Supplementary figure 2O-P), further validating that inhibitory inputs recorded following VR stimulation originated from Renshaw cells.

Our results show that dI3 neurons receive inhibitory inputs derived from efferent copies of motor outputs via direct inhibitory inputs from Renshaw cells (as well as excitation through sparse collaterals from motor neurons). Taken together, our observations suggest that dI3 neurons are well positioned to combine sensory feedback and efferent copies of motor outputs to mediate fast corrections of motor commands.

### dI3 neurons innervate motor neurons for flexion, extension, or co-activation

We next sought to quantify the projections of dI3 neurons to motor neurons: such projections – to the lateral motor column and to a lesser extent to medial motor neurons – have previously been described^18,20,43^. However, the identification of the motor neuron pools innervated was restricted to the pool innervating the quadriceps^20^. Therefore, we used Isl1^cre/wt^; Ai34d^fl/wt^ animals, restricting tdTomato expression to the presynaptic terminals of Isl1^ON^ neurons, combined with retrograde labelling of motor neurons innervating the ankle flexor, tibialis anterior (TA) or the ankle extensor, gastrocnemius (Gs, Figure 3A). We quantified tdTom^ON^/vGlut2^ON^ presynaptic boutons apposed to retrogradely labelled motor neurons; these terminals would not be from vGlut2^ON^ sensory neurons as the latter neurons project only to the dorsal horn^44,45^. We confirmed that motor neurons from the medial column in segments L4 and L5 receive only a few boutons from dI3 neurons, while motor neurons innervating both Gs and TA receive numerous dI3 terminals (Figure 3B,C; ^43^). Furthermore, the number of dI3 presynaptic terminals apposed to flexor motor neurons (TA) was significantly higher than those apposed to extensor motor neurons (Gs, 6.4 ± 1.6 and 4.7 ± 2.0 boutons per 100µm^2^ respectively, Figure 3C). Since the expression of vGlut2 by motor neurons has been reported but remains unclear^46,47^, we used Chat^cre/wt^; Ai34d^fl/wt^ animals to validate that presynaptic boutons quantified as dI3 terminals were not confounded with recurrent innervation between motor neurons^48^. We observed only a few tdTom^ON^/vGlut2^ON^ boutons from cholinergic neurons apposed to lateral motor neurons, which confirmed that the vast majority of terminals identified in the Isl1^cre/wt^; Ai34d^fl/wt^ animals were from dI3 neurons (Supplemental figure 3A-C).

**Figure 3:**
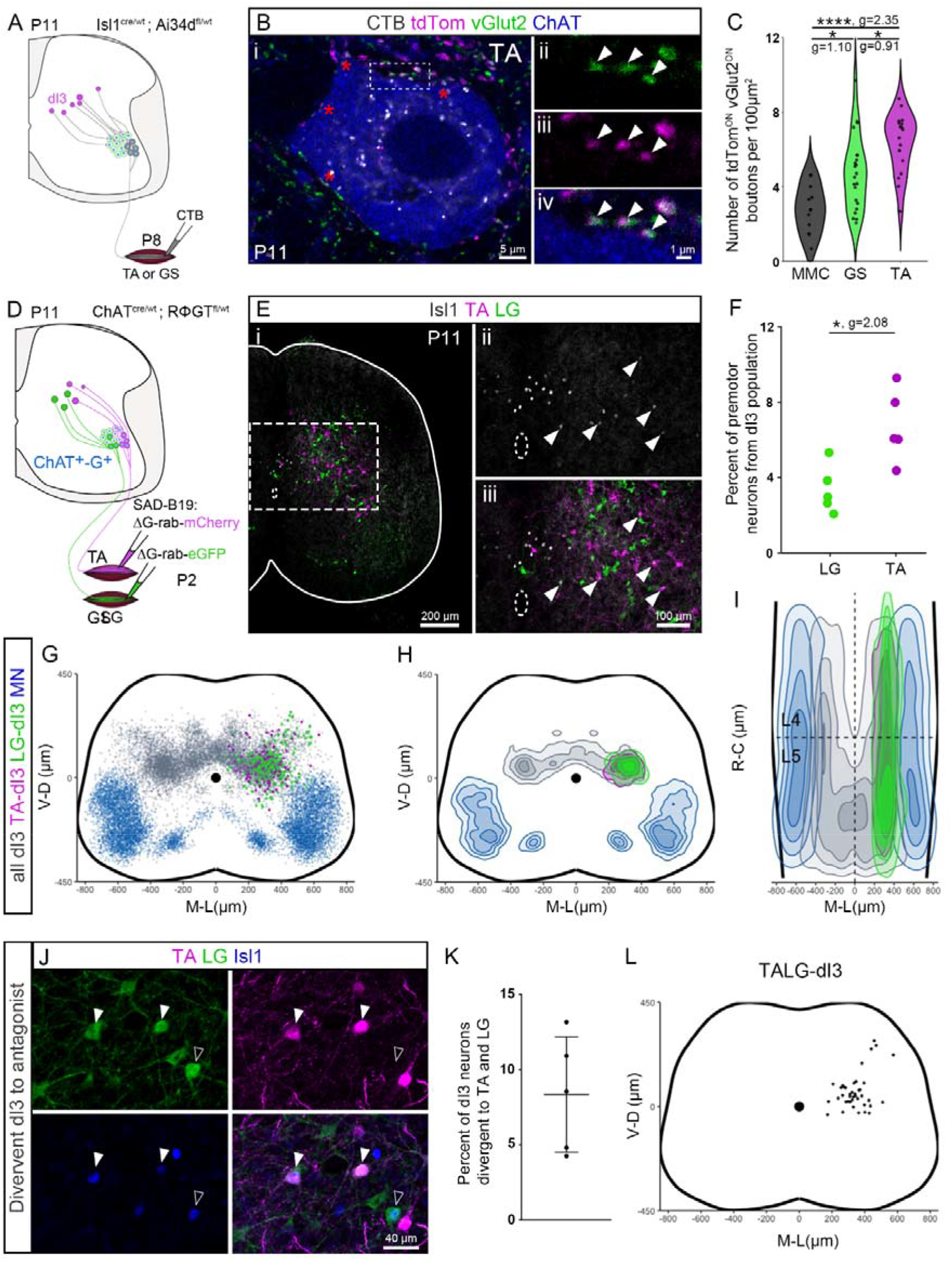
dI3 to motor neuron innervation is wired to mediate different types of motor output. (A) Experimental strategy to visualize presynaptic boutons from dI3 neurons (tdTom^ON^-vGlut2^ON^) apposing motor neurons innervating the tibialis anterior (TA) or the gastrocnemius (GS). CTB intramuscular injections were performed at P8 and the tissues collected at P11. (Bi) Representative example of a motor neuron innervating TA (CTB^ON^, in grey) with tdTom^ON^ (in magenta), vGlut2^ON^ (in green) double positive boutons apposed to its soma (ChAT, in blue). The double positive boutons apposed to the soma are highlighted with red asterisk or in the dashed box showed at higher magnification (ii-iv). (C) Number of tdTom^ON^-vGlut2^ON^ double positive presynaptic boutons apposed to the soma of motor neurons of the medial column or motor neurons innervating TA or GS scaled by their cross sectional area. (D) Monosynaptic retrograde tracing used to describe dI3 premotor neurons that project to motor neurons innervating the TA (SAD-B19-ΔG -mCherry) or the LG (SAD-B19-ΔG -GFP) with Rabies virus injection at P2 and tissue collection at P11. (Ei) Representative example of a lumbar transverse section showing premotor neurons traced from the injection in TA (mCherry, in magenta), LG (GFP in green) and a staining against Isl1 (in grey). (ii-iii) Part of the transverse section contained in the dashed box in (i) showing dI3 premotor neurons (Isl1^ON^, in grey) traced from motor neurons innervating the TA (Isl1^ON^-mCherry^ON^) or the LG (Isl1^ON^-GFP^ON^). White arrowheads highlight the premotor dI3 neurons. (F) Percent of premotor neurons innervating TA vs LG motor neurons that are identified as dI3 neurons (Isl1^ON^) scaled by the total number of premotor neurons traced from TA and LG injections respectively. (G-H) Spatial (G) distribution and (H) density of motor neurons (in blue), all dI3 neurons (Isl1^ON^ neurons, in grey) and dI3 neurons premotor to TA and LG motor neurons (respectively in magenta and green). (I) Spatial density showing the distribution of premotor dI3 neurons innervating TA and LG motor neurons along the rostro-caudal axis with a front view of the cord. (J) Orthogonal projection of a lumbar section following a premotor tracing from TA and LG, showing two dI3 neurons that are projecting to both TA and LG motor neurons (Isl1^ON^-mCherry^ON^-GFP^ON^, filled arrowheads) and one dI3 neurons innervating LG motor neurons (Isl1^ON^-mCherry^OFF^-GFP^ON^, contour arrowhead). (K) Percent of premotor dI3 neurons innervating both TA and LG motor neurons per animal. (L) Spatial distribution of dI3 neurons innervating both TA and LG motor neurons. (B-C) n=12 MMC motor neurons in N=6 animals, n=18 TA and 22 GS motor neurons in N=5 animals. (E-L) data shown are collected from L3 to L6, and pooled from N=5 animals. Non-hierarchically bootstrapped (C) Kruskal-Wallis test followed by Dunn’s multiple comparisons test and (F) two-tailed Mann Whitney test.

We then asked where the premotor dI3 neurons innervating flexor versus extensor motor pools are located, and whether some are wired for co-activation of flexor and extensor muscles. To tackle these questions, we used simultaneous monosynaptic retrograde rabies virus tracing from both extensor (the lateral gastrocnemius, LG) and flexor (TA) muscles. G-deleted rabies viruses were injected into LG and TA muscles of P2 pups expressing the G protein in cholinergic neurons (i.e. in motor but not sensory neurons, preventing anterograde tracing; Figure 3D, ^49,50^). Immunoreactivity against Isl1, visible in 40 to 50% of L3-L6 dI3 neurons at this age (Supplemental figure 3D-E) largely without spatial bias compared with the whole population of dI3 neurons (Supplemental figure 3F-H), was used to identify premotor dI3 neurons (Figure 3E). We found that the percentage of lumbar premotor neurons that are flexor-innervating dI3 neurons was greater than those innervating extensor motor neurons (6.7 ± 1.9% from TA vs 3.4 ± 1.3% from LG, Figure 3F), supporting the flexor bias observed using anterograde tracing (Figure 3C). Furthermore, premotor dI3 neurons were segregated to the interposed and most dorso-lateral region of the dI3 neuron population, with virtually no premotor dI3 neurons in the most medial subpopulation restricted to segments L5 and L6 of the lumbar enlargement (Figure 3G-H, Supplemental figure 3F-H, Supplemental Figure 1D). Although there was enrichment of premotor dI3 neurons to TA and LG in the same segments as the innervated motor pool (L4 and L5), premotor dI3 neurons were seen rostral to these segments as well (in L3), showing that some dI3 neurons project descending axons to at least the adjacent segment (Figure 3I, Supplemental figure 3K,^51^). Lastly, the simultaneous tracing from TA and LG showed that some dI3 neurons innervate both TA and LG motor neurons and are therefore positioned to mediate co-contraction of antagonist muscles (Figure 3J). Although the proportion of these divergent premotor dI3 neurons would be underestimated in these experiments (Figure 3K-L; ^52^), their proportion (∼8%) was relatively high compared to that which we previously described for the whole population of lumbar premotor neurons innervating TA and LG motor pools (∼4%; ^52^).

In summary, the position of premotor dI3 neurons spatially overlap with the location of dI3 neurons receiving multimodal sensory feedback and those receiving an efferent copy from Renshaw cells (and motor neurons). Furthermore, they are wired to contribute to flexion, extension, or co-contraction of antagonist muscles. Taken together, dI3 neurons, as a population, show key components of comparator modules.

### dI3 neurons form cellular comparator modules

We next sought to determine whether inputs from sensory afferents and efferent copies converge on the same premotor dI3 neurons. Our first step was to determine whether the muscles innervated by the motor neurons to which dI3 neurons project, supply proprioceptive inputs to the same dI3 neurons. To do so, we used intramuscular monosynaptic rabies tracing (same animals as Figure 3), which labels both premotor dI3 neurons and proprioceptive neurons innervating the injected and synergist muscles (Figure 4A, ^49^) but not afferents from antagonist muscles^53^. Although the proportions are largely underestimated due to the relatively low efficiency of monosynaptic retrograde tracing^52^, we nevertheless found that at least one third of premotor dI3 neurons also receive PV^ON^/vGlut1^ON^/RV^ON^ proprioceptive inputs from homonymous proprioceptive neurons (Figure 4B-E). Furthermore, this was the case when tracing from flexor as well as extensor muscles (Figure 4D, E).

**Figure 4:**
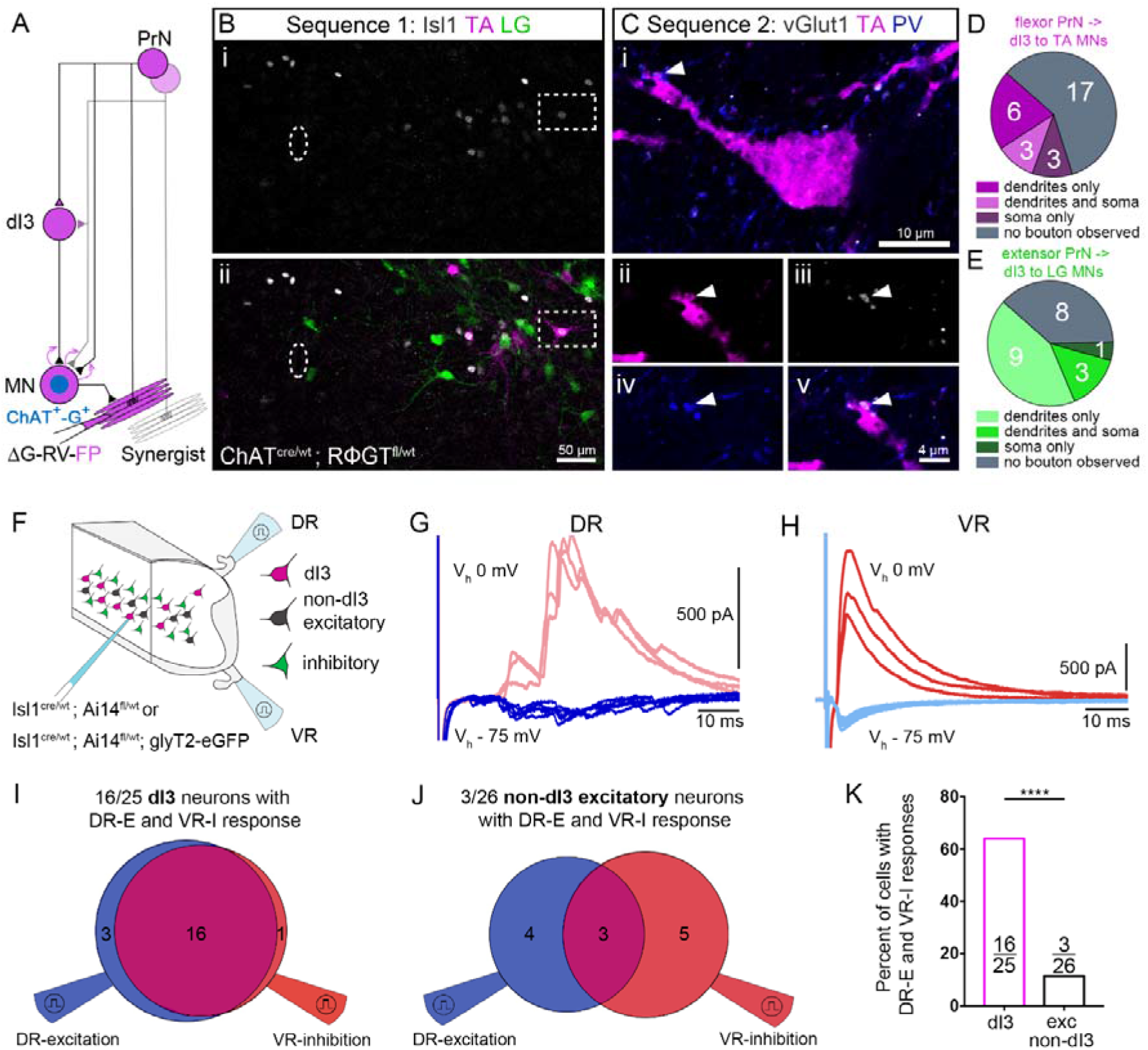
dI3 neuron circuits have connectivity of cellular comparator modules. (A) Following intramuscular injection of ΔG-RV in *Chat^cre/wt^; Rosa26^R^*^Φ*GT*^ mice, sensory and motor neurons innervating the injected muscles are primarily infected. The RV propagates to premotor neurons (here dI3 neurons), as well as proprioceptive neurons from targeted and synergist muscles. (B) Example of a representative lumbar transverse section after a first round of staining (sequence 1) used to identify premotor dI3 neurons, showing premotor neurons traced from the injection in TA (mCherry, in magenta), LG (GFP, in green) and anti-Isl1 (in grey). (C) The same section was then unmounted and reacted a second time (sequence 2) to visualize PV (in blue) and vGlut1 (in grey) and reimaged. After the second staining, the dI3 neuron premotor to TA motor neurons, highlighted in the dashed box in B, shows a presynaptic bouton from a proprioceptive neuron (vGlut1^ON^-PV^ON^) traced from the injection in TA (mCherry^ON^, in magenta). This shows that premotor dI3 neurons are receiving putative inputs from homonymous or heteronymous proprioceptive neurons. (D-E) Number of premotor dI3 to (D) TA and (E) LG motor neurons with apposed boutons from proprioceptive afferents of the homonymous or synergist muscles (vGlut1^ON^-PV^ON^-FP^ON^ bouton). (F) Schematic of the longitudinal preparation of the hemi lumbar cord used to record dorsal root (DR)-evoked as well as ventral root (VR)-evoked excitation and inhibition from dI3 neurons (tdTom^ON^, in magenta) or non-dI3 excitatory neurons (tdTom^OFF^GFP^OFF^, in black) in Isl1^cre/wt^;Ai14^fl/wt^ and Isl1^cre/wt^;Ai14^fl/wt^;glyT2-eGFP animals. (G-H) Representative traces for (G) DR-evoked and (H) VR-evoked responses recorded from a single dI3 neuron held at Vh - 75 mV for excitation and Vh 0 mV for inhibition. (I-J) Numbers of responsive (I) dI3 neurons and (J) non-dI3 excitatory neurons to DR-evoked excitation and VR-evoked inhibition shown as Venn diagrams. (K) Percentage of recorded neurons receiving both DR-evoked excitation and VR-evoked inhibition whether they are identified as dI3 or non-dI3 excitatory neurons (exc non-dI3). (B-E) N=2 animals traced from TA and LG. (G-K) Total of n=25 dI3 neurons from N=6 animals and n=26 non-dI3 excitatory neurons from N=3 animals (P5-14). (K) Fisher’s exact test.

We next assessed whether individual dI3 neurons receive convergent inputs derived from efferent copies (Renshaw cells) and sensory afferents. Following stimulation of dorsal as well as ventral roots on longitudinal hemisected spinal cord preparations (Figure 4F), we found that a large proportion of dI3 neurons receive both DR-evoked excitation and VR-evoked inhibition (16/25 recorded dI3 neurons from n=6 animals, Figure 4G-I). Ten of these also received VR-evoked excitation and DR-evoked inhibition (Figure 4G-H, Supplemental Figure 4A-B). This proportion is likely underestimated since in these preparations, some sensory axons are damaged. Indeed, the percentage of dI3 neurons with dorsal root evoked excitation in hemisected spinal cords is 20% lower (76%, 19/25 cells) than that seen in ventral spinal cord ablated preparations (97%, 36/37 cells), and the amplitude of the excitatory responses is one fifth (292 ± 337% vs 60 ± 57%, Figure 4G). To determine whether the convergence of these classes of inputs was specific to dI3 neurons or shared by other neuronal populations in the intermediate laminae, we recorded DR and VR-evoked responses in tdTom^OFF^/GFP^OFF^ non-dI3 excitatory neurons in preparations from animals expressing tdTomato in dI3 neurons and eGFP in inhibitory neurons (Figure 4F). In contrast to dI3 neurons, we observed that only a small proportion of tdTom^OFF^/GFP^OFF^ non-dI3 excitatory neurons receive both DR-evoked excitation and VR-evoked inhibition (3/26 recorded excitatory non-dI3 neurons from n=3 animals, Figure 4J-K), with 2/26 also receiving VR-evoked excitation and DR-evoked inhibition (Supplemental Figure 4C-D). That is, there was a high level of specificity of convergence of inputs from sensory afferents with those derived from efferent copies to dI3 neurons.

Together, these results suggest that a large proportion of dI3 neurons in the interposed region may form cellular comparator modules combining efferent signals and sensory feedback to modulate motor outputs.

### dI3 neurons mediate corrective abilities during complex motor tasks

We next sought to determine whether dI3 neurons play a fundamental role in correcting ongoing movements. To allow for reversible manipulation of dI3 neurons, the expression of the inhibitory DREADD receptor hM4D(Gi) was induced under the control of both Cre and Flp recombinases in Isl1^cre/wt^;vGlut2^flp/wt^ mice. The expression of the hM4D(Gi) receptor in Isl1^cre/wt^;vGlut2^flp/wt^ mice was induced by injecting AAV2/5-hSyn-Flp^ON^-DIO-HA-hM4D(Gi)-mCherry bilaterally along the lumbar spinal cord to restrict the manipulation to lumbar dI3 neurons (Figure 5A). We confirmed that hM4D expression was confined to Cre^ON^Flp^ON^ dI3 neurons and was not expressed in motor (given their expression of Isl1 and possible expression of vGlut2^46,47^) or sensory neurons (Figure 5B). We showed the effectiveness of expression in spinal cord slices: application of the agonist JHU37160 (230nM) led to a reduction of dI3 neuron excitability, seen as an increase in cell rheobase of ∼80%, together with a decrease in cell input resistance of ∼20%. Furthermore, the agonist had no effect on non-transduced, hM4D(Gi)^OFF^ dI3 neurons labelled with tdTomato, validating the specificity of its effect (Supplemental figure 5A-C).

**Figure 5:**
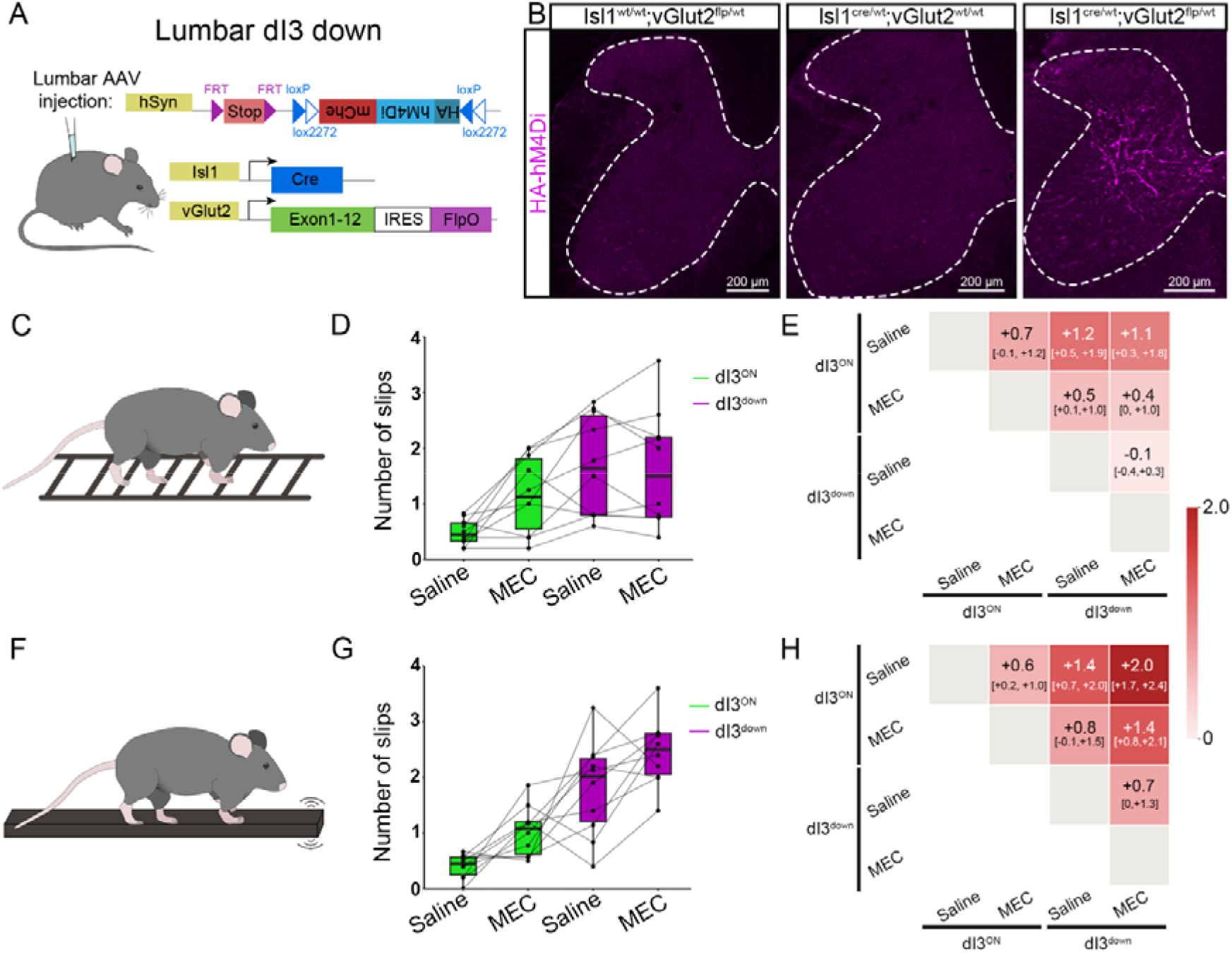
dI3 neurons mediate corrective responses using nicotinic signalling. (A) Intersectional strategies used to inhibit reversibly dI3 neurons activity combining genetic and viral transduction to specifically manipulate lumbar dI3 neurons. (B) Example of a representative lumbar transverse sections from Isl1^wt/wt^;vglut2^flp/wt^, Isl1^cre/wt^;vglut2^wt/wt^ and Isl1^cre/wt^;vglut2^flp/wt^ mice injected with a AAV2/5-hSyn-Flp^ON^-DIO-HA-hM4Di-mCherry demonstrating specific transduction of dI3 neurons only when both recombinases are expressed. (C, F) Schematics of the (C) uneven elevated horizontal ladder and (F) horizontal elevated vibrating beam tasks used to assess corrective responses. (D, G) Averaged number of hindlimb slips per mouse when crossing the elevated (D) ladder or (G) beam. Dots show the values for individual animals. dI3^ON^ (green) from a session preceded by a subcutaneous injection of saline together with an intraperitoneal injection of saline or mecamylamine (5mg/kg, MEC), while dI3^down^ sessions (magenta) were preceded by a subcutaneous injection of the DREADD agonist JHU37160. Values obtained from the same animal are connected with grey lines. (E, H) Matrix of the Hodges-Lehmann estimates of the within-animal differences in slips per crossing together with their 95% confidence intervals. Each cell gives the estimate of the column condition relative to the row with colour shading indicating the magnitude of the effect. (D-E, G-H) N=10 animals.

Since the presence of spinal cord modules combining sensory feedback and inputs derived from efferent copies could theoretically allow for detection of mismatches to generate corrective responses with minimal delays^3,9^, we next investigated the performance of mice during complex motor tasks. We turned to a behavioural test in which a horizontal ladder had unevenly spaced rungs. In the absence of hM4D(Gi), there were no differences in hindlimb slips when either saline or JHU37160 was given (0.4 ± 0.3 vs 0.3 ± 0.2 slips after saline and JHU37160 administration respectively, n=6 animals). In mice expressing hM4D(Gi) in lumbar dI3 neurons, however, the number of hind paw slips following JHU37160 administration increased (Figure 5C-E, Supplementary video 1), similar to what was previously described for a horizontal ladder test in a model of global dI3 output suppression^18^. Similar results were seen on an elevated vibrating beam (Figure 5F-H, Supplementary video 2).

To test whether there could be a role of efference copy from Renshaw cells in the corrective adjustments by dI3 neurons, we ran the same experiment preceded by an intraperitoneal injection of mecamylamine (5mg/kg), a nicotinic antagonist that disrupts Renshaw cell activity by inhibiting their activation by motor neurons (Supplementary figure 2L-P) but shows no significant effect on sensory inputs to dI3 neurons in longitudinal preparations (Supplemental figure 5D-F). In animals with no manipulation of dI3 neuron activity (dI3^ON^), mecamylamine administration led to an increase in the number of hindlimb slips compared to saline injected dI3^ON^ mice in both motor tasks (Figure 5D-E, G-H; Supplementary video 1). Conversely, when mecamylamine was administered to mice with lumbar dI3 neurons inhibited, it did not additionally increase the number of hindlimb slips on the horizontal ladder. These results suggest that its effects in dI3^ON^ animals could primarily have resulted from the reduction of Renshaw cell inputs to dI3 neurons on the ladder experiment (Figure 5D-E). On the more challenging vibrating beam walking, mecamylamine administration led to an increase in the number of slips (Figure 5G-H, Supplementary video 2), suggesting that in more challenging motor tasks, nicotinic transmission may be involved in corrective movements independently of dI3 neurons, for example via ventral spinocerebellar neurons^54^. Of note, in both tests, the number of slips in response to mecamylamine alone was lower than in mice with inhibited lumbar dI3 neurons (Figure 5D, E, G, H), which may have resulted from the incomplete block of efference copy (from Renshaw cells or another source), or may indicate that dI3 neurons also mediate fine corrective movements independently of the efferent copy they receive.

In summary, this set of experiments shows that dI3 neurons play a role in corrective motor adjustments during complex motor tasks. The results seen when inhibiting nicotinic receptors point to a potential role for Renshaw cells in the implementation of these corrections.

### dI3 neurons combine sensory feedback and efference copy to mediate reflexive corrections

To determine whether corrective responses that depend on an internal estimation of motor state are mediated by spinal circuits involving dI3 neurons, we turned to stumbling corrective reactions. Stumbling correction, originally described in cats and present in mice and humans, involves hyperflexion of the knee and the ankle in response to perturbation of the limb during swing phase, leading to stepping over the obstacle that would otherwise have prevented the progression of swing^12–14,55–57^. This reaction (i) is phase dependent, (ii) persists after lower thoracic spinal cord transection^13,58^, and (iii) is triggered within a few milliseconds^57^. Furthermore, given these features, stumbling correction, triggered by a sensory stimulus, (i) needs information about the phase of the step cycle, (ii) is generated locally within lumbar spinal cord circuits, and (iii) involves a minimal number of synapses. These three features led us to address whether the stumbling corrective response could be mediated by dI3 neurons.

To trigger a stumbling corrective reaction (SCR) in mice, we first applied a mechanical perturbation to the dorsal side of the paw during the swing phase of locomotion (Figure 6A)^14^. This stimulus resulted in a burst of muscle activity and hyperflexion of the ankle and knee (Figure 6B-C). We found that the maximum height of the step during a SCR was reduced following lumbar dI3 inhibition (Figure 6D, Supplemental figure 6B, Supplementary video 3). This reduced response was similar following administration of mecamylamine alone (Figure 6E, Supplementary video 3). Furthermore, when lumbar di3 neurons were inhibited, the inhibition of nicotinic transmission did not further reduce stumbling correction (Figure 6D-E, Supplemental figure 6B, Supplementary video 3). Taken together, these results suggest that dI3 neurons mediate stumbling corrective response using nicotinic-dependent signals and therefore suggest that dI3 neurons use Renshaw cell inputs to mediate these state-dependent reflexive corrections.

**Figure 6:**
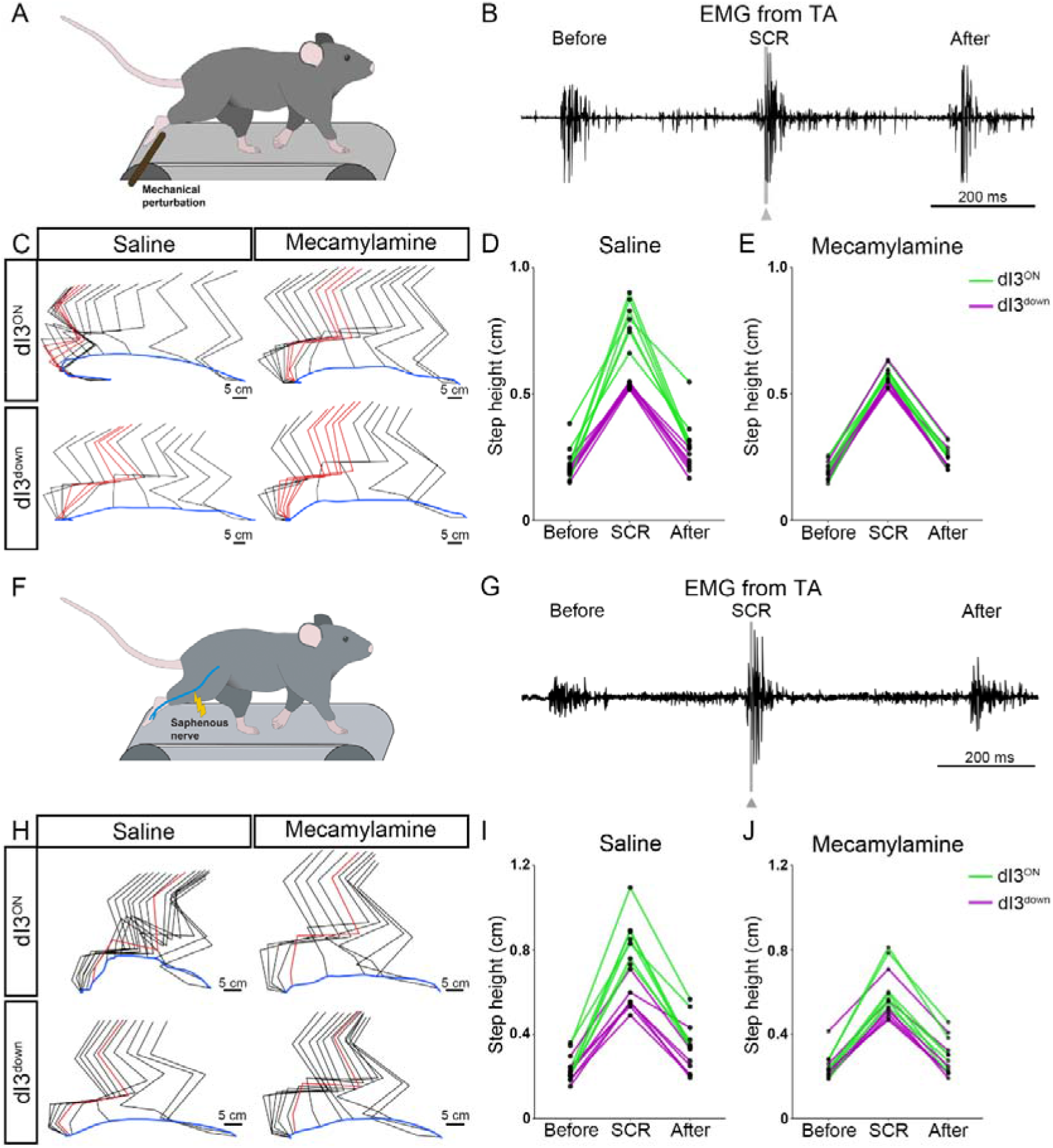
dI3 neurons modulate stumbling correction combining sensory feedback and nicotinic transmission, suggesting a role from Renshaw cell inputs. (A, F) Stumbling corrective reactions (SCR) were triggered using (A) mechanical perturbation or (F) electrical stimulation of the saphenous nerve. (B, G) Raw EMG of the tibialis anterior showing the step cycle before, during and after (B) mechanical or (G) electrical perturbation of the swing. The perturbation is indicated by the grey line and arrow on the EMG of the second swing. (C, H) Stick diagram reconstruction of a swing during the SCR with or without inhibition of dI3 neurons and with or without mecamylamine (5mg/kg). The SCR is triggered by (C) mechanical or (F) electrical perturbation. The time of the perturbation is shown in red and the trajectory of the toes in blue. (D, E) Step height of the step before, during and after mechanical SCR following intraperitoneal injection of (D) saline or (E) mecamylamine in animals without (dI3^ON^, green lines) or with (dI3^down^, magenta lines) inhibition of lumbar dI3 neurons. Lines connect averaged step height of the same animal, while dots represent averaged values of each animal. (I, J) Step height of the step before, during and after electrical SCR following intraperitoneal injection of (I) saline or (J) mecamylamine in animals without (dI3^ON^, green lines) or with an inhibition of lumbar dI3 neurons (dI3^down^, magenta lines). Lines connect averaged step height of the same animal, while dots represent averaged values of each animal. (D-E, I-J) N=7 animals.

In these experiments, there was a small increase in the height of the step preceding that with the mechanical perturbation in the dI3^ON^ animals injected with saline, raising the possibility that an anticipatory response might have interfered with the results (Figure 6D, Supplemental figure 6A). To circumvent this concern, we next induced a stumbling corrective response using electrical stimulation of the saphenous nerve during the early swing phase (Figure 6F). This stimulation reliably triggered a stumbling corrective reaction similar to that described previously^14^, with hyperflexion of the homonymous limb and a burst of activation in the ankle dorsiflexor (Figure 6G-H, Supplementary video 4). In these experiments, the height of the step before stimulation was similar in the different groups indicating a lack of anticipatory response (Figure 6I-J, Supplemental figure 6D). During SCR, we observed a decreased amplitude of the corrective reaction following lumbar dI3 inhibition (Figure 6I, Supplemental figure 6E, Supplementary video 4). Furthermore, step heights during SCR following mecamylamine administration alone or together with JHU37160 produced a similar reduction of reflexive correction (Figure 6I-J, Supplemental figure 6E, Supplementary video 4) – that is, as in the mechanical stimulation experiments, there was no additive effect. We therefore conclude that the reduction of the stumbling corrective reaction observed when dI3^ON^ animals were administered mecamylamine was due to the disruption of nicotinic-dependent circuits involving dI3 neurons, supporting a pivotal role of Renshaw cell inputs to dI3 neurons for the generation of reflexive corrections.

Interestingly, in the step succeeding both types of perturbation – mechanical or electrical – the effects of either JHU37160 or mecamylamine persisted, suggesting a potential role of this circuit in modulation over the longer term (Supplemental figure 6C, F).

Taken together, our results describe spinal circuits in which dI3 neurons combine the integration of sensory feedback with signals derived from efferent copies of motor outputs via Renshaw cells. We further show that this circuit is responsible for mediating state-dependent correction of ongoing movements by generating rapid adjustments.

## Discussion

In this study our aim was to determine whether spinal dI3 neurons form comparator modules for fast corrections of ongoing movements. Using circuit tracing, we describe the convergence of multimodal sensory inputs and inputs derived from efference copies, including via Renshaw cells, on dI3 neurons. We show that these two classes of inputs both project to individual dI3 neurons. In addition, dI3 neurons innervate motor neurons favouring flexor over extensor groups, with a subset forming divergent connections to antagonist sets of motor pools. By manipulating their activity during behaviour, we demonstrate that dI3 neurons play a pivotal role in corrective responses. We further show that nicotinic transmission is necessary for the generation of corrective dI3-dependent responses, suggesting that the input from Renshaw cells to dI3 neurons is essential for the generation of rapid corrections of movement. Based on this converging evidence, we thus propose that dI3 neurons are at the heart of spinal cord comparator modules (Figure 7).

**Figure 7:**
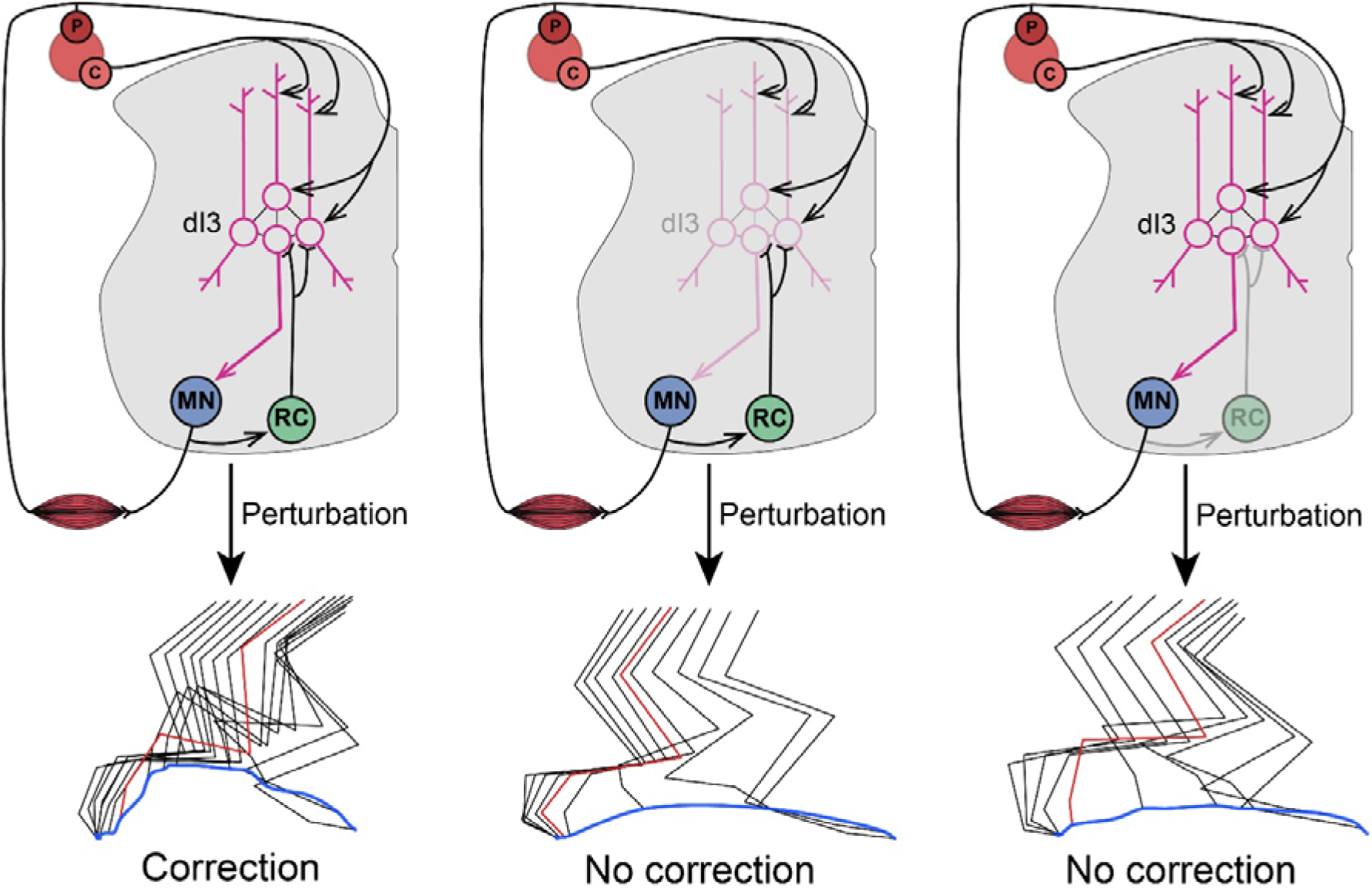
dI3 neurons combine sensory feedback and efference copies for corrections of ongoing movements. In intact circuits, dI3 neurons combine multimodal sensory feedback and efference copies mediated by Renshaw cells to induce fast correction of movements following a perturbation. When either dI3 neuron activity or nicotinic transmission are suppressed, fast corrections are reduced, suggesting the involvement of Renshaw cells in dI3 neuron circuits in the triggering of state-dependent reflexive correction of ongoing movements.

### Interposed dI3 neurons receive segregated, sensory inputs from multiple modalities

We have shown that dI3 neurons are widely distributed in the intermediate laminae, with positions differing along the rostro-caudal axis even within the lumbar spinal cord: for instance, the most medial dI3 neurons are seen primarily in the caudal segments. On this anatomical basis alone, it would therefore seem that this cardinal class, like others^59,60^, could form several subclasses that may differ molecularly, morphologically, and by their inputs, and thus may serve distinct functions.

It is clear that the position and the dendritic arborization of neurons are crucial to specify the inputs they receive^21,60–62^. We found that a large subpopulation of dI3 neurons located in an interposed position along the medio-lateral axis is distinct from others and itself likely comprises several sub-classes. This position overlaps with the “region of convergence” of proprioceptive axons from multiple muscles^21^, and these interposed dI3 neuronal somata are indeed densely innervated by proprioceptive afferents. Thus, the position of dI3 neurons may be critical to their function.

In contrast to these somatic proprioceptive inputs, cutaneous inputs are segregated to the dorsal dendrites of these neurons. This modality-dependent segregation of inputs across somato-dendritic compartments may have several important implications. Segregating inputs may confer a number of properties to neural circuits. For example, inputs to the distal dorsal dendrites of layer 5 pyramidal neurons in the primary somatosensory cortex allows for an increase in gain of tactile feedback during active perceptual detection^63,64^. Likewise, the spatial segregation of cutaneous inputs to dI3 neuronal dendrites may provide the substrate for specific modulation of these inputs^65^. Localizing cutaneous inputs to the dendrites may also provide a mechanism for specifically gating them, for example through proximal inhibitory inputs^66,67,68^, or amplifying them, for example through dendritic plateau potentials^69–71^.

This compartmental segregation may also allow more flexibility for the regulation of synaptic plasticity, a key component in the development of other comparator systems^72^. Given the fundamental role of dI3 neurons in plasticity of spinal circuits following complete transection^19^, these observations open new directions to further study the role of somato-dendritic compartmentalization in the integration of feedback and feedforward signals in spinal circuits.

### Renshaw cell inputs to dI3 neurons may convey expected reafferent input

To correct for perturbations whilst moving, three key pieces of data need to be taken into account by the nervous system: (a) the sensory feedback in response to the perturbation (exafference), (b) the sensory input resulting from the movement itself (“prediction” or reafference), and (c) the state of the motor output system, i.e. where on the input-output curves are motor neurons and motor pools. The latter is required to ensure that the corrective input to the motor pools leads to the intended motor output.

The stumbling corrective response, described in many species of limbed mammals, is ideal for studying corrective circuits, as it is a stereotypical, short-latency response that depends on the phase of the step cycle during which the stimulus is received^12–14,55^. While we have previously shown that dI3 neurons receive rhythmic inputs from spinal locomotor circuits^19^, we neither demonstrated the cellular substrate for these inputs nor suggested how they might “know” the input-output states of motor pools. Here, we use the stumbling corrective response to address these questions, and show that Renshaw cells are well situated to provide a source of locomotor efference copies to dI3 neurons. Furthermore, these inputs may also reflect the state of the motor pool^73^.

We note that our observation that a large proportion of dI3 neurons receives inputs from Renshaw cells may seem contradictory with previous studies that failed to find Renshaw cell-mediated inhibition of intermediate zone interneurons involved in reflex pathways from group I and II muscle afferents to motor neurons in cats^74^. In this region, dI3 neurons represent a small subset (<8%) of premotor neurons, and could not be specifically targeted in cats. Furthermore, we have found that some dI3 neurons have ascending projections^51^, and these would have been specifically excluded in the cat studies restricted to local interneurons. We conclude that the negative finding in cats cannot readily be extrapolated to dI3 neurons, although acknowledge that there may also be species differences.

Renshaw cell activity arguably represents the most direct and accurate negative efferent copy of motor commands^9^, as they are activated by motor neuron collaterals^15,75^. Since a negative (i.e. inhibitory) predictive signal is required to detect a mismatch between excitatory sensory feedback and the expected reafferent input^33^, we sought to acutely disrupt Renshaw cell activity. Previously, a genetic strategy was used to manipulate Renshaw cell activity and led to the suggestion that they may play a role in long term plasticity of spinal circuits^76^. We note, however, that the strategy used was not specific to Renshaw cells^39,76^ and in any case could not be used together with manipulation of dI3 neurons in the same animals. We therefore opted to use a pharmacological approach^42,77,78^, showing a reduction in the dI3-dependent response to perturbation by mecamylamine, which partially blocks motor neuron activation of Renshaw cells. Although we cannot exclude a concomitant effect on the modulation of other pathways, including primary afferents^79,80^, Ia inhibitory interneurons^81^ or ventral spinocerebellar tract neurons^54,82^, the evidence that the nicotinic effects were mediated via Renshaw cell inputs to dI3 neurons includes: (a) the synaptic connectivity, with Renshaw cells forming synapses with dI3 neurons (Figure 2A-K, Supplemental figure 2E-F); (b) the reduction in Renshaw cell activity with mecamylamine resulting in decreased inhibition of dI3 neurons (Supplementary figure 2L-P); (c) the minimal effects of mecamylamine on afferent transmission to dI3 neurons (Supplemental figure 5D-F); and (d) the use of this strategy in fictive preparations to show a reduction in Renshaw cell activation that resulted in reduced inhibition of motor neurons^42,77^. Thus, the preponderance of evidence points to Renshaw cell involvement in mediating the stumbling corrective response.

At first blush, it seems counterintuitive that disruption of an inhibitory negative image of the efference copy would lead to a *reduction* in the corrective response. Comparator neurons compare these inputs to external feedback signals – the removal of any of these inputs to comparators – whether positive or negative – would lead to a miscalculation of sensory prediction error and a failure to correct. This mechanism is in contrast to that of sensory gating^83^, which would lead to hyperflexion if the interneurons intercalated between the primary afferents and motor neurons were acutely disinhibited.

Mechanistically, the reduction in corrective response could result from the role of Renshaw cell inputs in the synchronisation of dI3 neuron firing. Inputs from Renshaw cells would necessarily precede afferent feedback and could therefore favour timely rebound activity upon this afferent input as they may do for motor neurons^84^, leading to increased flexion. Renshaw cells could also change the recruitment gain of the dI3 population^73^, ensuring their population activity matches that of motor pools. In addition, the microcircuit segregation of inputs to dI3 neurons that project to flexor vs extensor motor neurons is not known. Renshaw cells are known to inhibit the motor pool that activates them whilst disinhibiting antagonist pools via Ia inhibitory interneurons^85^. If there is a similar organisation with respect to dI3 neurons, it is possible that, in response to a stumbling stimulus during flexion, Renshaw cells inhibit extensor-related dI3 neurons and disinhibit flexor-related dI3s, in which case mecamylamine would reduce the corrective response. Furthermore, some responses may require co-contraction – thus it is conceivable that there are specific projections of Renshaw cells to sub-populations of dI3 neurons that innervate both flexor and extensor motor neurons.

In summary, there are several possible organizational and cellular mechanisms through which the absence of Renshaw cell input to comparators can contribute to a reduction of corrective responses. But in the absence of defined circuit diagrams – from motor neuron type to specific Renshaw cells to specific dI3 neurons and to classified motor neurons – suggestions such as these remain speculatory.

### A spinal comparator module

Our work suggests that Renshaw cells are a source of predictive information within spinal circuits, but predictive inputs are likely not restricted to this pathway. For instance, dI3 neurons – at least in the cervical spinal cord -also receive inputs from corticospinal neurons^86,87^. Descending inputs may modulate the corrective signals generated by dI3 neurons as has been suggested for spinal RORα neurons^28^. In fact, the effect of descending regulation of dI3 neurons may be reflected in the anticipatory response observed in the step prior to stumbling corrective reactions triggered by mechanical (i.e. visible) perturbation. Indeed, the step prior to the perturbation was significantly reduced when dI3 neurons were downregulated. Therefore, dI3 neurons may additionally process descending inputs to anticipate potential changes in the environment.

Comparisons between efference copies and instructive feedback occur in several brain circuits: such circuits have been extensively studied in cerebellar systems and in the neocortex^8,88,89^, and suggested for ventral spinocerebellar tract neurons^82^. However, the location of a comparator module in spinal circuits leads to an advantage in terms of minimising delays so as to trigger immediate corrections and prevent severe consequences of unexpected stimuli. Predictive processing in spinal circuits likely differs from the multifactorial tuning present in higher brain regions. Therefore, in addition to the generation of immediate corrections, dI3 neurons may relay these errors to higher centres^90^ to further refine corrections across longer time scales. Indeed, although corrections occur at different timescales and with different roles, circuits involved in the generation of predictive processing are not working in isolation. For instance, cerebellar outputs modulate neocortical predictive processing^88^, while climbing fibre activity associated with withdrawal reflexes may update cerebellar predictions following immediate corrections^91^. Therefore, in the future, it will be important to characterise how spinal reflexive corrections are used to update predictive signals in higher hierarchical centres.

## STAR Methods

### Animals

All experiments were performed according to the Animals (Scientific Procedures) Act UK (1986) and certified by the UCL AWERB committee, under project license number PP2688499 or under the approval of University of Ottawa’s animal care committee and conform to the guidelines put forth by the Canadian Council for Animal Care. Heterozygous *Isl1^cre/wt^* (Jackson Lab, stock #024242) were crossed with homozygous Ai14^fl/fl^ (Jackson Lab, stock #007914) or heterozygous Ai34d^fl/wt^ (Jackson Lab, stock #012570) to obtain respectively a somatic expression of tdTomato or its expression at presynaptic terminals. The same line was crossed with *Slc17a6*-IRES2-FlpO-D mice (referred as vglut2^flp/wt^, Jackson Lab, stock #030212) to allow the reduction of excitability in lumbar dI3 neurons. Heterozygous *En1^cre/wt^*(Jackson Lab, stock # 007916) were crossed with heterozygous Ai34d^fl/wt^ to induce tdTomato expression at the presynaptic compartment of V1 neurons. For experiments aimed at recording from Renshaw cells, heterozygous *Slc6A5e*^GFP/wt^ mice (termed glyT2-eGFP here, a gift from Prof. Zeilhofer, University of Zurich, ^92^) were used and further crossed with *Isl1^cre/wt^*Ai14^fl/wt^ to identify non-dI3 excitatory neurons. Homozygous *Chat^cre/cre^*mice (Jackson lab, stock #006410) were crossed with homozygous *Rosa26^R^*^Φ*GT*^ mice (Jackson Lab, stock #024708), to generate *Chat^cre/wt^; Rosa26^R^*^Φ*GT*^ that were used for rabies tracing experiments^50,52,93^. All mice used were in a C57BL6/J genetic background. For behavioural experiments, all mice were housed individually in a reverse light-cycle room with 12 hours of light from 7pm to 7am; all behavioural testing was performed in simulated darkness using red LED and infrared lighting between 8am and 5pm.

### Modified rabies virus production

The glycoprotein G-deleted variant of the SAD-B19 vaccine strain rabies virus was used (kind gift from Dr M.Tripodi). Modified RabV (ΔG-RV) with the sequence of the glycoprotein G replaced by the mCherry or eGFP sequence (ΔG-RV-eGFP/mCherry) was produced at a high concentration with minor modifications to the original protocol^94^. BHK cells expressing the rabies glycoprotein G (BHK-G cells) were plated in standard Dulbecco modified medium supplemented by 10 % foetal bovine serum (FBS) and split after 6–7hr incubating at 37°C and 5 % CO2. The cells were then inoculated at a multiplicity of infection of 0.2–0.3 with either ΔG-RV-eGFP or mCherry virus (initial samples kindly provided by Prof. Arber and Dr. Tripodi). Cells were incubated for 6 hours at 35°C and 3% CO_2_ and subsequently split 1 to 4 with 10% FBS medium and kept at 37°C and 5% CO_2_ for 12-24 hours. The medium was then replaced with 2% FBS supplemented medium and cells were incubated at 35°C and 3% CO_2_ for virus production. The supernatant was collected after ∼3 days and new medium was added for another round of production (three cycles maximum). The supernatant was filtered (0.45 μm filter) and centrifuged for 2 hours at 19 400 rpm (SW28 Beckman rotor). The pellets were suspended in PBS, dispersed and collected in a single tube and further centrifuged for 4 hours at 21 000 rpm in a 20% sucrose gradient (SW55 Beckman rotor). The resulting pellet was suspended in 100 μl PBS and the virus was stored at -80°C. The viral titre of each round of production was measured by serial 10-fold dilution of three different aliquots using standard protocols. The CSV-N2c rabies virus pseudotyped with EnvA used for the monosynaptic retrograde tracing from dI3 neurons were obtained from the Crick institute viral vector core.

### Intramuscular injections

Neonatal pups (P2 for SAD-B19-ΔG-RV injections, P7-8 for CTB injections) were anaesthetized using isoflurane inhalation and the skin above the injection site was wiped with ethanol 70%. For hindlimb injections, an incision was made on the skin to visualize the targeted muscle either tibialis anterior (TA), lateral gastrocnemius (LG, whose primary role is that of ankle extensors, but also contribute to knee flexion) or gastrocnemius (GS) without distinction between lateral and medial when injecting CTB. Each muscle was injected with 1 µL of SAD-B19-ΔG-RV or 0.5µL of CTB coupled to Alexa 488 (ThermoFisher, C34775) or 647 (ThermoFisher, C34778).

Subcutaneous injections in the hind paw were performed without incisions, using 0.5µL of CTB-Alexa 488 or Alexa 647.

The injected SAD-B19-ΔG-RVs were used at a titre between 10^9^ and 10^10^ IU/ml while CTB was used at 0.5%. All the injections were performed using a 5 μl Hamilton syringe (model 7652-01) fixed to a manual Narishige micromanipulator (M-3333) and loaded with a bevelled (30°) glass pipette of outer diameter 40-45 μm (1B100-4, World Precision Instrument). After slowly injecting the virus (> 1 minute), the needle was left on the injection site 2-3 minutes and then retracted. In the following 24 hours, injected pups were closely monitored to control for signs of movement impairment or rejection by the mother. The injected pups were perfused under terminal general anaesthesia respectively 9 days (SAD-B19-ΔG-RV) and 3-4 days (CTB) after injections.

### Intraspinal injections

AAVs to visualize dI3 somato-dendritic compartment were injected in P14-P16 animals while injections of helper AAVs were performed at 5-7 weeks old animals were injected to allow acute manipulation of lumbar dI3 neurons. Animals were first given Buprenorphine (Buprecare, XVD 130) to induce analgesia and then put under general anaesthesia with isoflurane (5% for induction, 1-1.5% maintenance). To stabilise the spinal cord of young pups a custom device for stabilisation of the vertebrae was used while adult animals were placed onto a stereotaxic frame (Kopf Instruments, Model 963 Ultra Precise Small Animal Stereotaxic Instrument). Intraspinal injections for anatomy and electrophysiology experiments were performed using a pulled and bevelled borosilicate glass pipette (Warner instruments, G120F-4) with an outer diameter of 25-35µm at the tip, attached to a stereotaxic device (Narishige, SMM-200). AAVs were injected applying multiple short pulses using a pneumatic picopump (World Precision Instruments, PV820). For behavioural experiments, injections were performed using a pulled and bevelled quartz capillary (Sutter Instruments, Q100-70-7.5) with tip at an outer diameter of 25-45 µm, attached to a Hamilton syringe (Hamilton, CAL7632-01) via a compression fitting (Hamilton, 55750-01), and mounted to the stereotaxic frame on a microinjector (Harvard Apparatus, Pump 11 Elite, Cat.# 70-4507). To label dI3 neurons prior to retrograde tracing from the sural nerve, a dorsal laminectomy of T13 was performed and 5 sites were injected on one side of the cord from L3 to caudal L5. For each site, 80 to 120nl of AAV1-phSyn1-flex-Lyn/EYFP-T2A-synaptophysin:mKate2 (final titer of 3.2 x 10^11^ vp/mL,VectorBuilder, ID VB180425-1066vcf, only the EYFP signal targeted by the Lyn domain to the membrane was used, abbreviated AAV1-phSyn1-flex-mYFP) was injected in the intermediate laminae (350µm medio-lateraly, 650µm dorso-ventrally) three weeks before tissue collection. To trace pre-synaptic neurons to dI3 neurons, about 75nl of a mix of AAV2/5-EF1a-flex-H2B-mTagBFP2-P2A-TVA (final titer of 1.8 x 10^12^ vp/mL, Viral vector core Crick Institute) and AAV2/5-Syn-flex-H2B-GFP-N2cG-CW3SL (final titer of 9.7 x 10^12^ vp/mL, Viral vector core Crick Institute), was injected in a single site in the intrathecal space between vertebrae T12 and T13 (300µm medio-lateraly, 350µm dorso-ventrally) in P21-24 animals. After 14 days, 75nl of CSV-N2c-EnvA-ΔG-mCherry was injected targeting the same location than for helper injection (final titer of 2.0 x 10^8^ vp/mL, Viral vector core Crick Institute) and the tissues were collected 7 seven days later. To induce the expression of the DREADD receptor hM4D(Gi) specifically in lumbar dI3 neurons, mice were injected with an AAV2/5-hSyn-Flp^ON^-DIO-HA-hM4D(Gi)-mCherry (final titer of 4 x 10^12^ vp/mL, 2064-aav5, Canadian Neurophotonics Platform Viral Vector Core Facility, RRID:SCR_016477). The intermediate laminae were injected on three sites on each side of the cord, 250-300nl per site, using the laminar spaces between T11-T12 (300µm medio-lateraly, 670µm dorso-ventrally), T12-T13 (350µm medio-lateraly, 700µm dorso-ventrally) and T13-L1 (350µm medio-lateraly, 650µm dorso-ventrally). To control the Cre dependence of each AAV construct, Cre negative animals were injected in the lumbar cord similarly to Cre positive animals, in at least 2 animals per construct and no fluorescent cells were observed at the sites of injection even after amplification of the fluorescent protein with immunoreaction. To control for the specificity of the monosynaptic retrograde rabies virus tracing from dI3 neurons, the same protocol was used on Cre negative animal and led to no labelling with mCherry in 2 animals. For this experiment, we also controlled for the presence of starter cells in the dorsal root ganglia following the tracing in *Isl1^cre/wt^* animals. We found only one DRG neuron expressing both helper construct as well as the mCherry from the rabies virus (Supplementary figure 2B). Since this sensory neuron was parvalbumin negative, it could not propagate the rabies virus anterogradely to motor neurons or Renshaw cells. To control the specific expression of double Cre and FlpO dependant AAV2/5-hSyn-Flp^ON^-DIO-HA-hM4D(Gi), the construct was injected in Cre^ON^-Flp^OFF^ and Cre^OFF^-Flp^ON^ animals. A maximum of up to 5 cells were labelled in the sections adjacent to the injection sites (45 μm thickness), with none detected further than 90 μm from the injection site. These represent a negligible proportion of labelled cells in Cre^ON^-Flp^ON^ animals used for behavioural experiments

### Retrograde labelling from the sural nerve

To label the cutaneous sensory axons from the sural nerve, P30 mice previously injected with AAV1-phSyn1-flex-mYFP were anaesthetized using isoflurane following induction of analgesia. An incision was made above the distal limb and the nerve was isolated from the facia and cut. The nerve was then put on a parafilm, surrounded by vaseline and dipped into a solution of CTB-Alexa 647 at 0.5% (Thermo Fisher Scientific, C34778) for 1 hour. The extremity of the nerve was washed multiple times with saline and the overlying skin was sutured without reattachment of the extremity of the nerve. The animals were then perfused five days after the procedure. After perfusion, the dorsal root ganglions of L3 to L6 were collected and stained for parvalbumin and CTB to verify the identity of sensory neurons labelled. Slices selected randomly were imaged, and 8/197 (n=2 animals) sensory neurons quantified expressed parvalbumin, confirming that the vast majority of sensory boutons traced were from cutaneous sensory neurons.

### Patch-clamp electrophysiology

#### Spinal cord preparations and solutions

Mice were first anaesthetized with a mixture of ketamine/xylazine (100 mg/kg and 10 mg/kg, respectively) and decapitated. The spinal column was dissected and a ventral laminectomy was performed in a chamber filled with cold (∼2°C) artificial cerebrospinal fluid (aCSF, the same used for recordings) composed of (in mM): 113 NaCl, 3 KCl, 25 NaHCO_3_, 1 NaH_2_PO_4_, 2 CaCl_2_, 2 MgCl_2_, and 11 D-glucose continuously bubbled with 95% O_2_ and 5% CO_2_. To map the functional inputs to dI3 neurons several slicing approaches were used to stimulate ventral and/or dorsal L4-L5 roots while recording from dI3 neurons, motor neurons or Renshaw cells. After laminectomy, the lumbosacral cord was glued to an agar block (7% agar/0.1% methyl blue) either ventral or dorsal horn facing up for experiments requiring ventral and dorsal-root stimulations, respectively. The tissue was then placed in a chamber filled with ice-cold slicing aCSF (∼2°C) comprising (in mM): 130 K-gluconate, 15 KCl, 0.05 EGTA, 20 HEPES, 25 d-glucose, 3 Na-kynurenic acid, 2 Na-pyruvate, 3 Myo-inositol, 1 Na-l-ascorbate, pH 7.4 with NaOH^95^ and sliced longitudinally^96^ or with an angle (∼45°, 350 µm thickness) using a vibratome (HM 650 V, Microm, Thermo Fisher Scientific). Slice(s) were incubated in a chamber with recording aCSF (constantly bubbled with 95/5% O_2_/CO_2_) at 37°C for about 30min, then maintained and recorded at room temperature (∼22°C). For hemisected longitudinal preparation, we did not slice the cord using a vibratome, but instead carefully hemisected the cord in recording aCSF before performing the recordings.

To investigate synaptic inputs in response to ventral or dorsal root stimulation (see^96^ for further details) in voltage-clamp mode, a caesium-based intracellular solution containing QX-314 to block spikes was used. This intracellular solution included (in mM): 125 Cs-gluconate, 4 NaCl, 0.5 CaCl_2_, 5 EGTA, 10 HEPES, 3 QX-314-Br, 2 Mg-ATP, pH 7.3 with CsOH, and osmolarity of 290–310 mosmol. Inhibition and excitation were recorded at their calculated equilibrium potential (+15 and -60 mV, taking into account the calculated junction potential of ∼-15 mV for all the intracellular solutions we used). To assess the inhibitory effect of DREADD agonist JHU37160, we used GTP-enriched, ATP-regenerating intracellular solution containing (in mM): 140 K-gluconate, 10 KCl, 0.1 EGTA, 10 HEPES, 2 Mg-ATP, 0.5 5′-GTP-Na_2_, 5 phosphocreatine-Na_2_, pH 7.3 with KOH, and osmolarity of 290–310 mosmol. For loose-cell attached recordings the electrode was filled with recording aCSF.

#### Recording configurations and analysis of responses

Ventral- or dorsal-horn ablated longitudinal spinal cord preparations from Isl1^cre/wt^;Ai14^fl/wt^ were used to infer about excitatory and inhibitory drive from sensory afferent or motor efferent activation to dI3 neurons. We used borosilicate glass pipettes (∼1.5x the root diameter) to stimulate dorsal or ventral L4 and/or L5 roots. We initially patched a motor neuron to define the stimulus threshold, i.e. the minimum stimulus intensity that generates a clear excitatory synaptic response, and then used 2-3x threshold intensity for the rest of the experiment. Following root stimulations (an average of 10 sweeps with interstimulus interval of ∼10 seconds and stimulus duration of 0.1-0.2 ms), the stimulus artifact was subtracted from the recording trace with a single or double exponential, and baseline-to-peak amplitude (size) and the artifact-to-first response time (latency) for each peak were calculated. Root evoked excitatory (EPSC) or inhibitory postsynaptic currents (IPSC) were measured in voltage clamp, holding the cell at the reversal potential for each ion (0 mV for cations and -75 mV for anions). Taking into account the junction potential of -15 mV, this corresponds to a dialled holding potential of -60 mV and +15 mV for EPSCs and IPSCs respectively. The synaptic conductances were calculated by dividing the observed currents by the driving force for cations and anions. In the case of multipeaked responses, we calculated the size and the latency of the first peak. The amplitude and latency of the EPSCs and IPSCs were averaged across sweeps, whereas the jitter was calculated as the variance of the latency for all sweeps. For dI3 neurons which responded to root stimulations, cell coordinates along mediolateral axis were taken. In these set of experiments, we also injected a 500ms-long current to calculate i) the membrane time constant, ii) cell conductance and iii) capacitance in current clamp mode^97^.

In the subset of dorsal-horn ablated preparations (on the last cell patched in the preparation), stimulation of the ventral root was repeated after addition of 3 µM 2,3-dihydroxy-6-nitro-7-sulfamoyl-benzo(f)quinoxaline (NBQX, Hello Bio, HB0442) and 50 µM 2-amino-5-phosphonopentanoic acid (APV or AP5, Hello Bio, HB0225) to block AMPA and NMDA receptors respectively.

Recordings of Renshaw cells were performed in oblique slices obtained from glyT2-eGFP animals. First, we recorded from a lumbar motor neuron to set the stimulation intensity for ventral root stimulation as previously described. Next, we identified Renshaw cells based on their i) location (laminas VII and IX), ii) expression of eGFP and iii) presence of extracellular spikes in response to ventral root stimulation in loose-cell attached mode. We performed whole-cell recordings from motor neurons and loose-cell-attached recordings from Renshaw cells, to measure ventral root-evoked extracellular spikes in Renshaw cells and IPSCs in motor neurons over a minimum of 10 sweeps. We then perfused 100µM mecamylamine (Sigma, M9020) for at least 15 minutes. Additionally, in spinal cord slices obtained from Isl1^cre/wt^;Ai14^fl/wt^ mice, we recorded ventral root-evoked inhibitory and dorsal root-evoked excitatory responses from dI3 neurons before and after application of 100 µM mecamylamine.

Hemisected preparations from In Isl1^cre/wt^;Ai14^fl/wt^ animals were used to measure ventral and dorsal root responses from dI3 neurons. We stimulated ventral and dorsal roots (L4 or L5 roots) with the stimulation intensity set as 2-3x the threshold required to elicit an initial synaptic response in dI3 cells measured in whole-cell mode. In Isl1^cre/wt^;Ai14^fl/wt^;glyT2-eGFP animals, we conducted the same experiments to additionally record from non-dI3 excitatory neurons. We first performed patch-clamp recordings from dI3 neurons to determine the threshold stimulation intensity and confirm the quality of the preparation, and then we targeted eGFP^OFF^/tdTomato^OFF^, i.e. non-dI3 excitatory neuron. For these sets of experiments, we excluded 2 hemisected preparations in which there were no cells showing a response to stimulation of the dorsal root, suggesting that most of the sensory axons had been damaged. To validate the effect of the DREADD receptor agonist JHU37160 on silencing dI3 neurons expressing hM4D(Gi), we performed whole-cell recordings from dI3 neurons in oblique spinal cord slices from adult Isl1^cre/wt^;vglut2^flp/wt^ mice. These mice were injected with AAV2/5-hSyn-Flp^ON^-DIO-HA-hM4D(Gi)-mCherry and age-matched to those used in behavioural experiments. To assess the specificity of JHU37160, we also recorded from dI3 neurons in non-injected adult Isl1^cre/wt^;Ai14^fl/wt^ mice. We estimated cell input resistance using a 500ms-long current step and determined rheobase as the minimum current eliciting a spike during the ascending phase of a triangular ramp protocol (4 s up, 4 s down). These measurements were conducted before and after applying 230 nM JHU37160, repeated every 5 minutes for at least 15 minutes.

#### Equipment

Cells were observed using a Nikon Eclipse E600FN microscope (Nikon, Japan) equipped with a dual-port system for simultaneous imaging of infrared differential interference contrast (DIC) via a digital camera (Moment-mono-OC) and fluorescence. Excitation was provided by a 488nm LED for EGFP expressing cells, and a 560nm LED for cells expressing TdTomato (Opto LED, Cairns Instruments, UK), and emission was captured using a CCD camera (Retiga XR, QImaging, UK).

Whole-cell recordings were performed using an Axopatch 200B amplifier (Molecular Devices, Sunnyvale), loose-cell attached recordings were done using an ELC-03X amplifier (NPI Electronics), and signals were filtered at 5 kHz and acquired at 50 kHz using a Digidata 1440 A A/D board (Molecular Devices, Sunnyvale) and acquired using Clampex 10 software (Molecular Devices, Sunnyvale). Patch pipettes were pulled from borosilicate glass (GC150F, Harvard Apparatus, Cambridge, UK) with a Flaming-Brown puller (P1000, Sutter Instruments, CA) to a resistance of ∼5 MΩ for dI3 neuron whole-cell and Renshaw cell loose-cell attached recordings, and ∼2 MΩ electrodes were used for motor neuron recordings. The roots were tightly sealed with glass micropipettes cut to similar diameters of L4 and L5 roots, and then used to stimulate the axons through an isolated constant current stimulator (DS3, Digitimer, Welwyn Garden City, UK).

### EMG electrode and nerve cuff fabrication

To record muscle activity and electrically stimulate afferent fibres in the hindlimb during treadmill walking, custom EMG muscle and nerve cuff electrodes were made following the protocols described in previous mouse studies^14,98,99^. Briefly, for muscle implant electrodes, stainless steel wire (AM System, #793200) was inserted into the head piece connector (Samtec, #CLP-108-02-L-D) and attached to a 27G needle on the remaining end. The nerve cuff electrode was fabricated with the same stainless steel wire, connected to the same head piece while the end of the electrode wires was inserted through a silicone cuff (Implantech, PAT02) to be slipped around the saphenous nerve and tied shut with a silk suture running through the silicone cuff (Ethicon BV-1, K809H). The plastic housing of the connector was then coated in a thin layer of biocompatible gel Epoxy (opaque 5-Minute Epoxy Gel, Devcon 14265). After allowing the Epoxy coating to cure completely for at least 24 hours, implants were then sterilized prior to surgery in 3% hydrogen peroxide solution for 20 minutes followed by a sterile water wash to remove any residual hydrogen peroxide.

### Nerve cuff and EMG electrode implantation

Following recovery after intraspinal injection, surgical implantation of EMG electrodes and nerve cuff was performed. Animals were first given Buprenorphine to induce analgesia and then put under general anaesthesia with isoflurane (5% for induction, 1-1.5% maintenance). EMG electrodes were implanted into the tibialis anterior and lateral gastrocnemius, of both right and left hindlimbs. Additionally, one nerve stimulation cuff electrode was placed around the saphenous nerve of the left hindlimb. Electrodes were implanted following the surgical procedures previously described^99,100^. Briefly, incisions were made in the upper neck region and in the skin of the hindlimbs to expose the target muscles and saphenous nerve. The electrodes were drawn through the neck incision down to the target muscle or nerve regions and inserted into the muscle parallel to the fibres or placed around the saphenous nerve. A head piece connector was secured into the skin of the dorsal neck region to allow stimulation from external equipment. Mice were given at least 1 week to recover before any acclimation or training sessions began prior to behavioural testing.

### Tissue collection and immunohistochemistry

The mice were perfused with PBS (0.1 M) followed by PBS 4 % paraformaldehyde (PFA, ThermoFisher, 28908) under terminal ketamine/xylazine anaesthesia (i.p. 80 and 10 mg/kg, respectively). Spinal cords were collected and post-fixed for 2-3 hr in 4% PFA, cryoprotected in 30 % sucrose in 0.1M PBS, embedded in PolyFreeze (Polysciences, 25113-1) or OCT (Tissue-Tek, 4583) and sliced with a cryostat (Leica, CM3050 S).

To visualize cutaneous boutons from the sural nerve and dI3 neurons, tissues were perfused from P35 animals, the spinal cord was sectioned in 100 µm thick free-floating sections were first blocked 30 minutes in PBS 2-X (0.2 M), 0.2-0.3 % Triton 100-X, 10% donkey normal serum, and then incubated with primary antibodies for 72 hours at 4°C and with secondary antibodies 24h at 4°C in the same blocking solution.

To quantify the number of dI3 presynaptic boutons onto motor neurons from P11 mice, 50 µm thick sections on slides were first treated with citrate buffer 0.3% (ThermoFisher, 036439.A3), pH 6, at 80°C for 3 minutes. Then sections were blocked 30 minutes in PBS 2-X (0.2 M), 0.2 % Triton 100-X (Sigma, T9284), 10 % donkey normal serum (Sigma, D9663), and incubated with primary antibodies for 72 hr at 4°C and with secondary antibodies 24 hr at 4°C in the same blocking solution.

For all the other immunohistochemistry, sections on slides (30-40 µm thickness) were made from P11 mice to visualize proprioceptive axons and localize premotor dI3 neurons using monosynaptic retrograde tracing from TA and GS; and from P6-7 animals to observe calbindin positive V1 terminals on dI3 neurons. They were first blocked 30 minutes in PBS 2- X (0.2 M), 0.2-0.3 % Triton 100-X, 10 % donkey normal serum, and then incubated with primary antibodies for 32-36 hours at 4°C and with secondary antibodies overnight at 4°C in the same blocking solution. To assess the presence of homonymous/heteronymous proprioceptive boutons onto premotor dI3 neurons following premotor tracing with RV, sequential immunoreaction and imaging were performed. The first reaction was performed on GFP, mCherry, parvalbumin (PV) and Isl1 as described above. Then, the tissues were imaged, the slides unmounted and incubated with primary antibody against vGlut1 and PV for 24 hr followed by an incubation overnight at 4°C with secondary antibodies.

The primary antibodies used were: goat anti-choline acetyltransferase (ChAT, 1:100, Millipore, AB144P), rabbit anti-CTB (1:400, Abcam, ab34992), goat anti-CTB (1:5000, List Labs, 703), chicken anti-mCherry (1:2000, Abcam, Ab205402), rat anti-RFP (1:1000, Chromotek, clone 5F8), chicken anti-GFP (1:1000, Abcam, Ab13970), rabbit anti-GFP (1:2500, Abcam, Ab290), goat anti-GFP (1:500, Abcam, ab6673), guinea pig anti-GFP (1:750, Synaptic systems, 132 005), rabbit anti-Calbindin (1:1000 to 1:4000, Swant, CB38a), rabbit anti-PV (from 1:1000 to 1:2000, Swant, PV27a), guinea pig anti-vGlut1 (1:2000, Millipore, AB5905), guinea pig anti-vGlut2 (1:2000, Millipore, AB2251-I) and guinea pig anti-Isl1 (1:7500, RRID:AB_2801512 from Susan Brenner-Morton, Columbia University, New York); and the secondary antibodies: donkey anti-goat Alexa 647 (1:1000, ThermoFisher, A-21447), donkey anti-goat Alexa 488 (1:1000, ThermoFisher, A11055), donkey anti-goat preabsorbed Alexa 405 (1:200, Abcam, Ab175665), donkey anti-guinea pig Alexa 647 (1:700, Millipore, AP193SA6 and 1:1000, Jackson ImmunoResearch, 706-605-148), donkey anti-guinea pig Alexa 488 (1:500, Jackson ImmunoResearch, 706-545-148), donkey anti-guinea pig DyLight 405 (1:400, Jackson ImmunoResearch, 706-475-148), donkey anti-rabbit Alexa 488 (1:1000, Thermo Fisher, A21206), donkey anti-rabbit Alexa 405 (1:500, Abcam, 175649), donkey anti-chicken Cy3 (1:1000, Jackson ImmunoResearch, #703-165-155), donkey anti-chicken CF 488A (1:500, Sigma, SAB4600031), donkey anti-rat Alexa 555 (1:500 to 1:1000, Thermo Fisher, A78945). In some experiments, Neurotrace Blue (1:200, ThermoFisher, N21479) or Alpaca anti-mTagBFP-647 (1:100, Synaptic System, N0501-AF647-L) were used together with the secondary antibodies to visualize the shape of neurons somata or BFP expression when needed. Free floating sections were mounted in Ce3D tissue clearing solution (BioLegend, 427704) while sections on slides were mounted in Mowiol (Sigma, 81381) and coverslipped (VWR, #631-0147) for imaging.

### Pharmaceutical intervention prior to behaviour testing

To reduce dI3 neurons activity, JHU37160 (dihydrochloride, DREADD ligand, Hello Bio, HB6261) was administered subcutaneously at a dosage of 0.5mg/kg. For dI3^ON^ groups, the equivalent dosage/volume was administered as saline subcutaneously. To modulate cholinergic neurotransmission, mecamylamine (hydrochloride, Cayman Chemical, 826-39-1), was administered intraperitoneally at a dosage of 5mg/kg to effectively reduce Renshaw cell activation, while the equivalent volume of saline was injected for saline condition^42,101^. The mecamylamine treatment, was used in combination with either JHU37160 or saline to create 4 treatment conditions, listed as follows: dI3^ON^ + saline, dI3^ON^ + mecamylamine, dI3^down^ + saline and dI3^down^ + mecamylamine. All the behavioural tests were performed in a time interval between 15 and 45 minutes after drug administration. All mice performed behavioural sessions for the 4 treatment conditions. These sessions were separated by at least 48 hours to allow for complete drug metabolism and to prevent *hM4D(Gi)* receptor desensitization.

### Training regimen and behavioural testing

Following recovery after electrode implantation surgeries, all mice were given 2 weeks (total of 10 sessions) of training and acclimation to the testing apparatuses and behavioural testing room. The first week consisted of strictly treadmill training. The second week combined both treadmill training and acclimation of the other testing apparatuses. For acclimation to the other apparatus, mice were placed on a horizontal ladder with all ladder rungs in place and evenly spaced and then on the balance beam, without vibration.

Following training and acclimation (∼4-5 weeks after intraspinal injections), 9-12 weeks old mice were tested on three different behavioural tasks. For treadmill trials, mice were placed on a horizontal treadmill set to 0.04m/s and walked for approximately 5 minutes. For half of the total treadmill sessions, electrical perturbations were triggered at the onset of the swing phase via electrical stimulation of the saphenous nerve. Treadmill trials were then repeated for the remaining half of behavioural sessions with mechanical perturbation with the use of a thin metal rod to physically perturb the swing of the hindlimb during walking. Electrical stimulation of the saphenous nerve was performed using live EMG recordings of the TA muscle from a custom closed-loop Axoscope protocol (Molecular Devices), wherein threshold detection of bursting activity from the TA muscle triggered an electrical pulse sent to the saphenous nerve cuff implanted in the left hindlimb (0.2ms duration, 100Hz, ranging from 0.1-0.3mA). Stimulus intensity of the pulse to the saphenous nerve was determined individually for each mouse as 1.2 times the required current to elicit a small response from the TA muscle at rest. Throughout the duration of treadmill trials, muscle activity from TA and LG hindlimb muscles were recorded simultaneously with Axoscope software (Molecular Devices, 10 kHz sampling frequency). For each condition, on each mouse, stumbling corrective reaction was triggered at least twice by saphenous nerve stimulation and twice by mechanical perturbation. The maximum step heights was then averaged per mouse for each condition.

Mice were then tested on a horizontal ladder and balance beam tests, both 60 cm long. In each testing session, mice performed on average 5 attempts with a successful crossing of the ladder and beam. To increase the difficulty of these tasks, the horizontal ladder was set up with unevenly spaced rungs and the balance beam, 0.9cm wide, was equipped with a coin flat vibrating motor (12000rpm, Blinli Amazon Shop). The horizontal ladder rungs were uniquely rearranged prior to each testing session to prevent a learning effect. All videos were recorded at 100 fps for stumbling corrective reactions, horizontal ladder and beam walking (Basler, ace U acA640-750um area scan camera, Cat.# 106748). Number of slips reported correspond to the average number of slips to cross the beam/ladder.

#### Imaging and analysis

All images were obtained using a Zeiss LSM800 confocal microscope. Images of the hemisections or of the entire coronal sections were obtained using x10 or x20 air objectives (0.3 and 0.8 NA respectively). High magnification images to visualize and quantify presynaptic boutons were taken using a 63x oil objective (1.4 NA). Tiles were stitched using Zen Blue (ZEN Blue 2.3 software) and analyses were performed using Imaris (Bitplane, version 9.6.1) and ImajeJ (version, 1.54k) softwares.

All the anatomical analysis on dI3 neurons were performed systematically from L2 to L5 for the monosynaptic retrograde rabies virus tracing from dI3 neurons and from L3 to L6 for all the other experiments. The quantification of presynaptic boutons was performed manually, following the creation of a colocalization channel for PV/vGlut1. Since Isl1 is also expressed in sensory but not corticospinal neurons, PV^OFF^/vGlut1^ON^/tdTom^ON^ could be identified as putative cutaneous boutons. A colocalization channel was also created for Calb/tdTom or vGlut2/tdTom to analyse putative Renshaw cell and dI3 boutons onto motor neurons respectively. In each case, the colocalization channel was used to facilitate the visualisation of boutons of interest and the expression of individual markers was subsequently confirmed on every bouton quantified. Cell mappings of premotor dI3 neurons following ΔG-RV injections were manually performed on every other section, in order to minimize the risk of counting the same cell twice in two consecutive sections. Putative Renshaw cells and motor neurons presynaptic to dI3 neurons, were mapped on every other slices when Calbindin positive cells at the ventral edge of the grey matter were labelled with mCherry, The cells that were calbindin and mCherry positive in a different domain of the cord were not included as they could belong to another cell population. Starter dI3 neurons (GFP^ON^BFP^ON^mCherry^ON^) were mapped on every section to control that no motor neurons were potential starter cells in the experiment as they also express Isl1. Other analyses were carried out on one every 4 slices from L3 to L6 to assess the rostro-caudal variation of the quantified parameters. Distribution maps were plotted setting the central canal as (0,0) in the (x,y) Cartesian system and using the ‘Spots’ function of Imaris. The y-axis was set to the dorso-ventral axis. Positive values were assigned for dorsal neurons in the y-axis and ipsilateral neurons in the x-axis. Coordinates were normalized through the cervical and lumbar parts separately using white mater borders and fixing the width and the height of the transverse hemisections^50^. The normalization of coordinates was performed independently for each quadrant using the following reference points: the x dimension was normalized to the outer edge of the white matter at the level of the central canal, while the y dimension was normalized for each quadrant using the outermost points of the white matter. The resulting cylindrical reconstruction of the spinal cord was then scaled to an idealized spinal cord size for illustrational purposes. All coordinate scaling and mapping were performed using a custom script in R adapted to read.csv files generated by Imaris (R Foundation for Statistical Computing, Vienna, Austria, 2005, http://www.r-project.org, version 3.6.2).

When presynaptic boutons on dI3 neurons were imaged, the hemislices were first acquired with a x10 or x20 air objective, and the position of dI3 neurons taken at high magnification were pinpointed on the hemislice to recover their localisation following quantification performed on high magnification images (x63 objective). To observe homonymous/heteronymous proprioceptive boutons onto premotor dI3 neurons following RV injections, the ipsilateral hemislices were first imaged with a x20 objective after staining Isl1, GFP and mCherry and subsequently imaged at high magnification with a x63 oil objective after the additional reaction of PV and vGlut1 to quantify the boutons on previously pinpointed premotor Isl1 positive neurons. The first step of imaging allowed to identify premotor dI3 and reimaged them during the second round of imaging based on their position and morphology, with occasionally the Isl1 staining that was still visible.

Following monosynaptic retrograde tracing from lumbar dI3 neurons, the dorsal root ganglions of the spinal segments injected were collected, all the slices were imaged and the expression of helper constructs as well as mCherry was checked on all slices to confirm the absence of proprioceptive (PV^ON^ GFP^ON^BFP^ON^mCherry^ON^) neurons that could act as starter cells.

All video recordings were analysed with DeepLabCut Python software^102^ (Version: 2.3.10) to label and track the joints of the hindlimb, and to extract automated hindlimb coordinate data to assess stepping and gait kinematics. Pose estimation data from DeepLabCut was then analysed with the ALMA toolbox Python software^103^ (Automated Limb Motion Analysis; Version: 1.1.0) to score the number of paw slips for each attempt across the horizontal ladder and balance beam, as well as to extract various kinematic parameters of stepping on the treadmill.

### Statistical analysis

Statistical analysis were performed using RStudio (v2024.4.2.764). The difference between groups were considered significant for *p* < 0.05. The statistical tests used are define in the figure legend and the level of significance are reported as follow: \**p* < 0.05, \*\**p* < 0.01, \*\*\**p* < 0.001, \*\*\*\**p* < 0.0001, ns. indicates no significance. In the text, summary data are reported as mean ± SD.

The data produced using electrophysiological or anatomical methods have an intrinsically nested structure, in which observations are taken from the cells, in a specific animal including sometimes an intermediate level, the root stimulated during *in vitro* recordings. Therefore, to analyse the data collected, we first identified the origin of the variance in our datasets. The interclass correlation coefficient (ICC) was calculated using a one-way random effect ICC model (1,1) with the ICCbare function of the ICC R package. When ICC≤0.5 in all the groups to be compared, we treated the data as independent and applied all the following tests to the lowest level of observation, i.e. the cell. When ICC≥0.5 in at least one of the groups, we performed a hierarchical bootstrap. We first resampled with replacement the animals (first level) and then within each animal resampled the cells with replacement (second level)^50,104^. Since the ICC between the root stimulated (L4 and L5) and the cell levels was very low in all our datasets, this hierarchical level was not taken into account in our analysis.

After ICC calculation, the normality of the distributions was tested using a Shapiro-Wilk tests. The groups compared from electrophysiological datasets were considered as paired for all the parameters as the responses were measured for excitation and inhibition in the same recorded cell following the stimulation of the same pool of axons (ventral or dorsal L4 and L5 roots). For one parameter, when a measurement was possible solely for the inhibitory or the excitatory response following a root stimulation, the cell was excluded from the analysis and the plots shown as there was no pairing possible. For anatomical datasets, the groups were independent. Accordingly, the bootstrap replicas were computed with paired and non-paired resampling and the Hedges’s G coefficient with (g_rm) or without pairing using the R function cohens.d from the “effsize” package. Where mean differences are reported, it corresponds to the distribution of 5 000-10 000 bootstrap replicas, the mean and the 95% CI of the distribution.

For behavioural experiments, animals were used as experimental units. Because normality of the distribution could not be assessed with sufficient power, we used distribution free methods, with effects reported as estimates with confidence intervals. The effects are calculated as the Hodges-Lehmann estimator (HL(x) = median((x_i_+x_j_)/2) for 1 ≤i ≤j ≤n). For the beam and ladder experiment, the estimator was applied to the within-animal differences with effects reported in number of slips. For the SCR experiments, it was applied to the within-animal log ratios and the result back-transformed, such that effects are reported as percent changes (calculated as 100 x (exp(HL of the log ratios) - 1)). The 95% confidence intervals were obtained by inverting the signed-rank test: the interval is bounded by the k^th^ smallest and k^th^ largest Walsh average, where k is set by the exact null distribution of the signed-rank statistic, such that it comprises every null value the test would not reject at the 5% level. Intervals are per comparison and unadjusted for multiplicity.

## Resource availability

### Material availability

All materials used are publicly available, except plasmids of AAV constructs used for monosynaptic retrograde rabies tracing. These constructs are available upon request to the lead contact or directly to the vector viral core of the Francis Crick Institute.

## Acknowledgements

We would like to thank Elzbieta Jankowska for thorough and helpful discussions of this work as well as for planning of future experiments. We thank Susan Morton for providing the Isl1 antibody, Kathryn Moreland and Ben Holahan for assistance with behavioural testing and Julie Tremblay for her assistance with animal care. We thank Molly Strom and the viral vector core of the Crick Institute for providing the helper vectors and the CSV-N2c rabies virus for tracing experiment. This work was supported by a Royal Society Newton International Fellowship (NIF\R1\192316) to M.G.O.; a Sir Henry Wellcome Postdoctoral Fellowship (221610/Z/20/Z) to F.N.; Canadian Institute of Health Research Project Grants (PJT 180556) to T.V.B. and (PJT 162357) to T.A.; a Wellcome Trust Discovery Award (227433/Z/23/Z) to R.M.B. and M.B.; a Biotechnology and Biological Sciences Research Council Research Grant (BB/S005943/1) to M.B. and a Wellcome Trust Early Career Award (225674/Z/22/Z) to R.R. R.M.B. is supported by Brain Research UK.

## Declaration of interests

R.M.B. is a co-founder of Sania Therapeutics Inc.

---

**Figure Supplemental 1:**
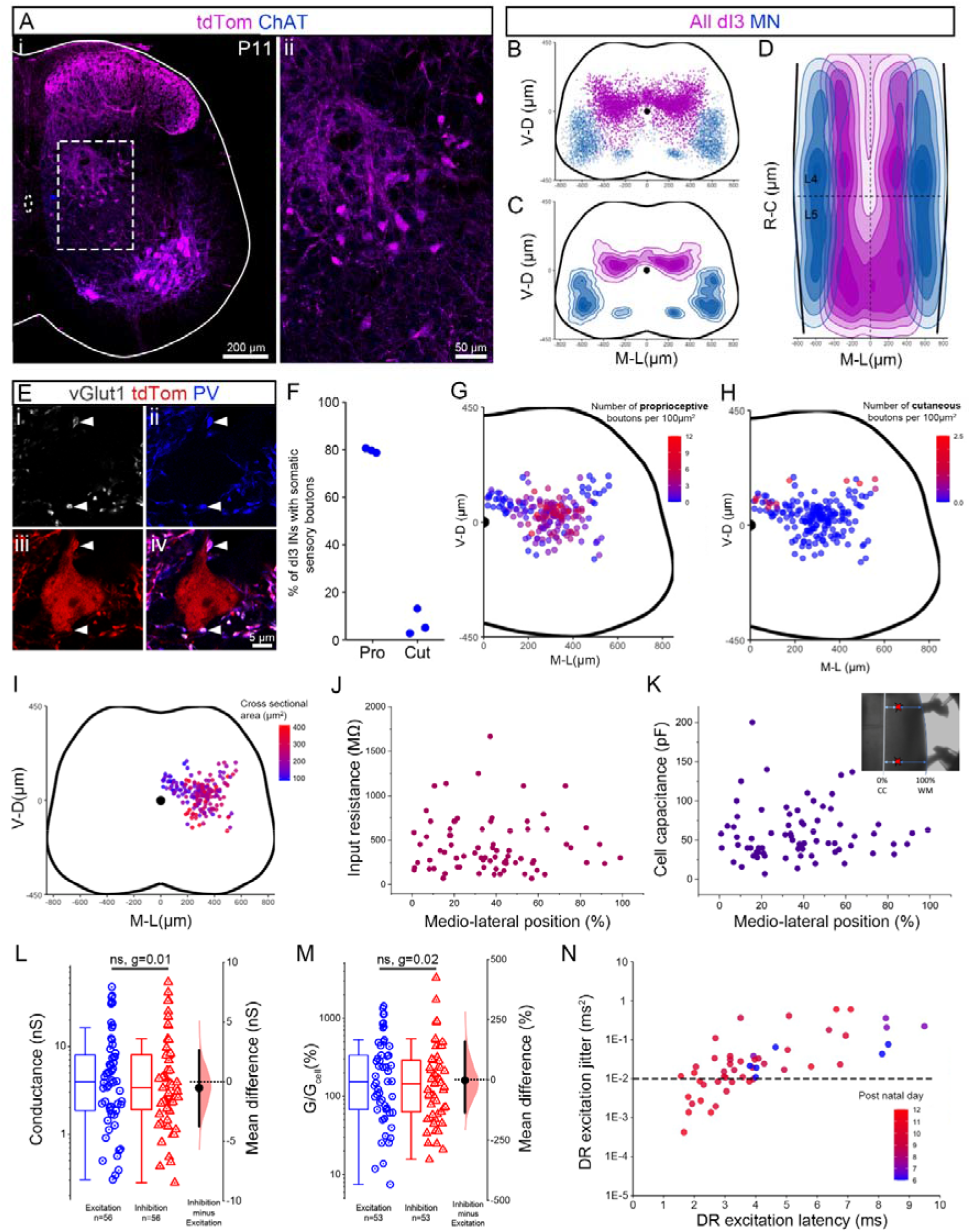
dI3 neurons segregate spatially based on sensory inputs they receive but not based on their passive properties. (A) Representative example of a lumbar transverse hemisection at P11 showing dI3 neurons (tdTom^ON^ChAT^OFF^, in magenta^ON^blue^OFF^). (ii) Part of the transverse section contained in the dashed box in (i) showing dI3 neurons (tdTom^ON^ChAT^OFF^) at a higher magnification. (B-C) Spatial (B) distribution and (C) density of all dI3 neurons (in magenta) and motor neurons (in blue) on an idealized lumbar transverse section. (D) Spatial density showing the distribution of the same population as in B-C along the rostro-caudal axis with a horizontal view of the cord from L3 to L6. (E) Example of a dI3 neuron (tdTom^ON^) with proprioceptive boutons (vGlut1^ON^-PV^ON^-tdTom^ON^) apposed to its soma. (F) Percentage of lumbar dI3 neurons with somatic proprioceptive (Pro, vGlut1^ON^-PV^ON^-tdTom^ON^) vs cutaneous (Cut, vGlut1^ON^-PV^OFF^-tdTom^ON^) boutons apposed to their soma. (G-H) Spatial distribution of dI3 neurons colour coded by the number of (G) proprioceptive or (H) cutaneous boutons apposed to their soma scaled by their cross sectional area. (I) Spatial distribution of dI3 neurons colour coded by the cross section of their soma. (J-K) Medio-lateral distribution of (J) input resistance and (K) cell capacitance of dI3 neurons along the medio-lateral axis. Inset in (K) describes the coordinate system used to measure the medio-lateral position of recorded neurons. (L-M) Comparison of the size of the DR-evoked excitation and inhibition (L) raw and (M) scaled to cell conductance. Box plots are accompanied by the mean difference on the right, n shows the number of plotted root responses from L4 and L5. (N) DR-evoked excitation jitter vs latency colour coded by the age of the animal used. The dashed line indicates a conservative estimate of the cut-off for monosynaptic window based on Doyle and Andresen (2001), as determined in adult brain stem. (A-I) The same N=3 animals were used as in figure 1A-E. (A-D) Data shown are from L3 to L6 pooled from the 3 animals at P11, (E-I) n =192 dI3 neurons analysed. (J-K) Recording obtained in (J) n=77 and (K) n=73 dI3 neurons from the same preparation as in figure 1J-O and figure 2J-K. (L-N) Recordings obtained from the same N=5 animals, 2 roots and n=37 dI3 neurons as in figure 1J-O. Plotted values in L and M are restricted to root stimulations that triggered both excitatory and inhibitory measurable responses in the recorded dI3 neurons to allow hierarchically bootstrapped paired samples Wilcoxon tests and paired Hedge’s G.

**Figure Supplemental 2:**
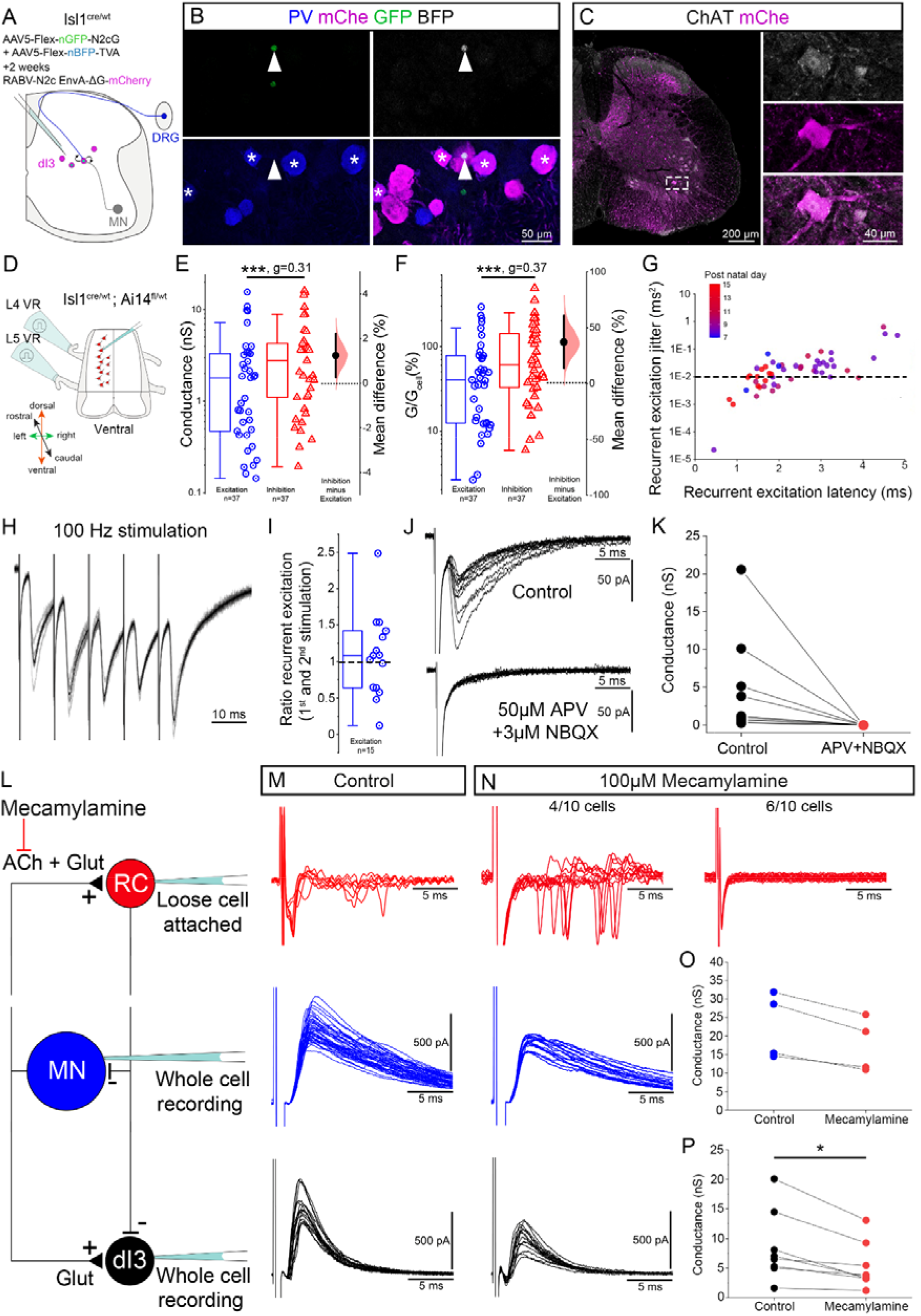
dI3 neurons receive monosynaptic purely glutamatergic inputs from motor neurons. (A) Schematic of the method used to trace presynaptic partners to dI3 neurons. The two helper AAVs were injected in the lumbar cord of juvenile Isl1^cre/wt^ mice followed two weeks later by the injection of CSV-N2c-EnvA-ΔG-mChe. (B) Image of a section of a dorsal root ganglion (DRG), showing one of the two potential starter DRG neuron (mChe^ON^GFP^ON^BFP^ON^, white arrowhead) found after our tracing. Since both DRG neurons were negative for PV, it cannot lead to an anterograde labelling of a motor neuron or a Renshaw cell. Multiple DRG neurons are presynaptic to dI3 neurons (mChe^ON^) with some proprioceptive (PV^ON^, white asterisk) and mechanosensory (PV^OFF^) neurons innervating to dI3 neurons. (C) Example of a lumbar transverse hemisection showing a mChe^ON^ChAT^ON^ motor neuron (dashed box, at higher magnification on the right side). (D) Schematic of the preparation used to record ventral root (VR)-evoked responses in dI3 neurons. (E-F) Comparison of the size of recurrent inhibition and excitation conductance, (E) raw and (F) scaled to cell conductance where n is the number of root stimulations (L4 and L5) recorded. (G) Plot indicating recurrent excitation jitter vs latency. Dashed line indicating a conservative estimate of the cut-off for monosynaptic window based on Doyle and Andresen (2001), as determined in adult brain stem. (H) Representative trace of recurrent excitation following VR high frequency stimulation at 100Hz. (I) Ratio of the amplitude of recurrent excitation measured between the response to the first and second VR stimulation at 100 Hz. (J) Representative traces for recurrent excitation in control conditions (top) and in the presence of glutamate receptor antagonists (bottom). (K) Size of the recurrent excitation in control and glutamate-blocked conditions. (L) Schematic representing the connectivity of Renshaw cells, motor neurons and dI3 neurons, together with the type of recording configuration used to test the effect of mecamylamine on VR-stimulation responses. (M) Sample responses obtained from a Renshaw cell (red, loose cell attached recording), motor neuron (blue, whole-cell recording) and dI3 neuron (black, whole-cell recording) after ventral root stimulation in control condition. (N) Same responses recorded after 100 µM mecamylamine application into the recording solution. Example Renshaw cell in top right is the same neuron illustrated in (M), as are the motor neuron and dI3 neuron. (O-P) Comparisons of the recurrent inhibition size in control vs 100 µM mecamylamine conditions recorded from (O) motor neurons and (P) dI3 neurons. (E-F) Recordings obtained from the same 8 mice (P5-15), 2 roots and 40 dI3 neurons as in Figure 2J-K. In E and F, plotted values are restricted to root stimulations that triggered both excitatory and inhibitory measurable responses in the recorded dI3 neurons to allow non-hierarchically bootstrapped paired samples Wilcoxon tests and paired Hedge’s G. (I) n= 15 dI3 neurons recorded from N=3 animals at P7. (K) n=10 root stimulations in N=5 animals (P7-15) and n=6 dI3 neurons.(M-P) n= 10 Renshaw cells were recorded from N=4 glyT2-eGFP animals, while n=6 dI3 neurons and n=4 motor neurons were recorded in a total of N=3 and 4 Isl1^cre/wt^;Ai14^fl/wt^ animals respectively. (P) Non-hierarchically bootstrapped paired samples Wilcoxon tests.

**Figure Supplemental 3:**
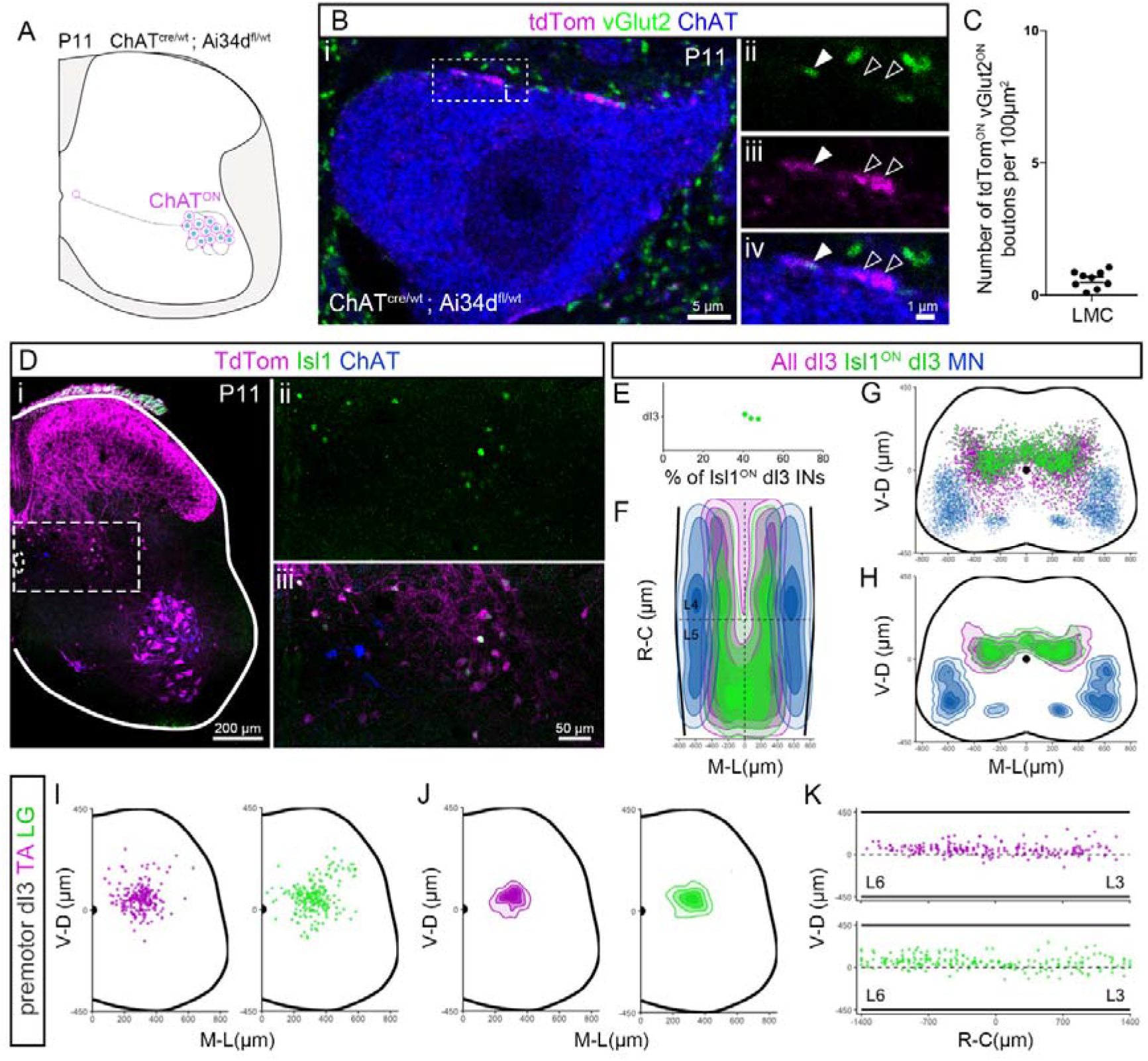
Most dI3 innervation of motor neurons arises from interposed di3 neurons. (A) Representative example of a motor neuron (ChAT, in blue) with one tdTom (in magenta), vGlut2 (in green) double positive bouton apposed to its soma in a ChAT^cre/wt^; Ai34d^fl/wt^ mouse at P11. The double positive bouton apposed to the soma is highlighted with the dashed box in i, shown at higher magnification in ii-iv. Filled arrowhead shows the tdTom^ON^vGlut2^ON^ bouton and contour arrowhead the tdTom^ON^vGlut2^OFF^ boutons. (C) Number of tdTom^ON^-vGlut2^ON^ double positive presynaptic boutons apposed to the soma of motor neurons from the lateral motor column scaled by their cross-sectional area. (D) Representative example of a lumbar transverse hemisection from a Isl1^cre/wt^; Ai14^fl/wt^ mouse at P11 showing dI3 neurons (tdTom^ON^ChAT^OFF^, in magenta) stained with an antibody against Isl1 (in green). (ii-iii) Part of the transverse section contained in the dashed box in (i) showing dI3 premotor neurons (tdTom^ON^ChAT^OFF^) at higher magnification, with about half of them still expressing Isl1 (in green). (E) Percentage of dI3 neurons (tdTom^ON^ChAT^OFF^) labelled by immunostaining on Isl1 (tdTom^ON^ChAT^OFF^Isl1^ON^) at P11 in the lumbar cord. (F) Spatial density showing the distribution of all dI3 (in magenta), dI3 neurons immunoreactive for Isl1 (in green) and motor neurons (in blue) from L3 to L6 in a horizontal view of the cord. (G-H) Spatial (G) distribution and (H) density of the same population as in (F) on an idealized cross section of the lumbar spinal cord. (I-J) Spatial (I) distribution and (J) density of ipsilateral premotor dI3 neurons innervating TA (in magenta) and LG (in green) on an idealized hemi-section. (K) Spatial distribution of dI3 neurons premotor to TA and LG along the rostro-caudal axis with a horizontal view of the lumbar spinal cord. (C) n=9 LMC motor neurons in N=3 animals. (D-H) Data shown are from L3 to L6 pooled from N=3 animals and collected on the same samples as in Figure Supplemental 1A-D. (I-K) Same dataset as Figure 3E-L, N=5 animals pooled.

**Supplemental Figure 4:**
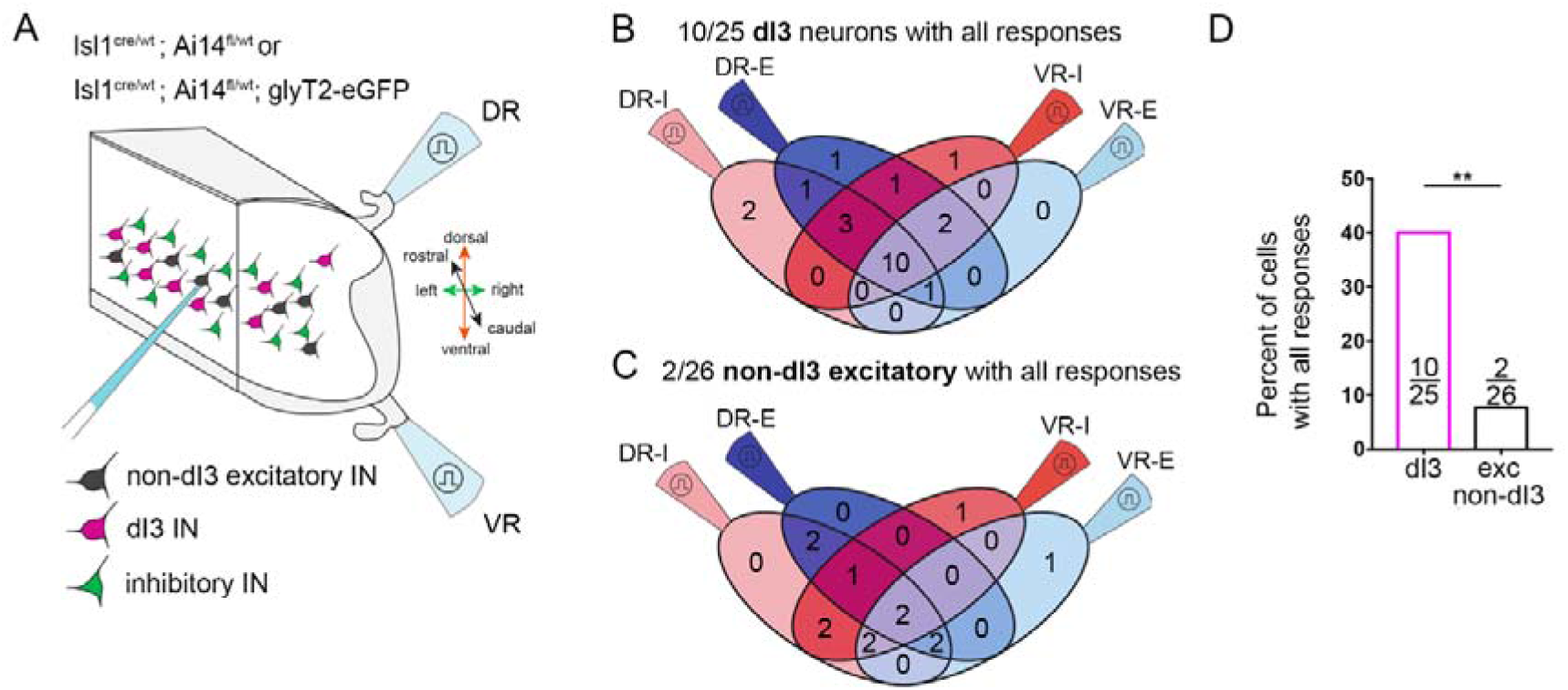
dI3 neurons in the intermediate laminae receive a specific set of inputs. (A) Schematic of the lumbar hemicord preparation (as in Figure 4F) from Isl1^cre/wt^;Ai14^fl/wt^;glyT2-eGFP or Isl1^cre/wt^;Ai14^fl/wt^ animals used to record (DR)-evoked as well as (VR)-evoked excitation and inhibition from dI3 neurons (tdTom^ON^, in magenta) or non-dI3 excitatory neurons (tdTomato^OFF^GFP^OFF^, in black). (B-C) Distribution of (B) dI3 neurons and (C) non-dI3 excitatory neurons depending on their responses to VR and DR-evoked stimulations. (D) Percentage of recorded neurons receiving DR and VR-evoked excitation and inhibition whether they are identified as dI3 or non-dI3 excitatory neurons (exc non-dI3). (B-D) Same n=25 dI3 neurons and n=26 non-dI3 excitatory neurons from respectively N=6 and 3 animals (P5-14) presented in figure 4I-K. (D) Fisher’s exact test.

**Supplemental Figure 5:**
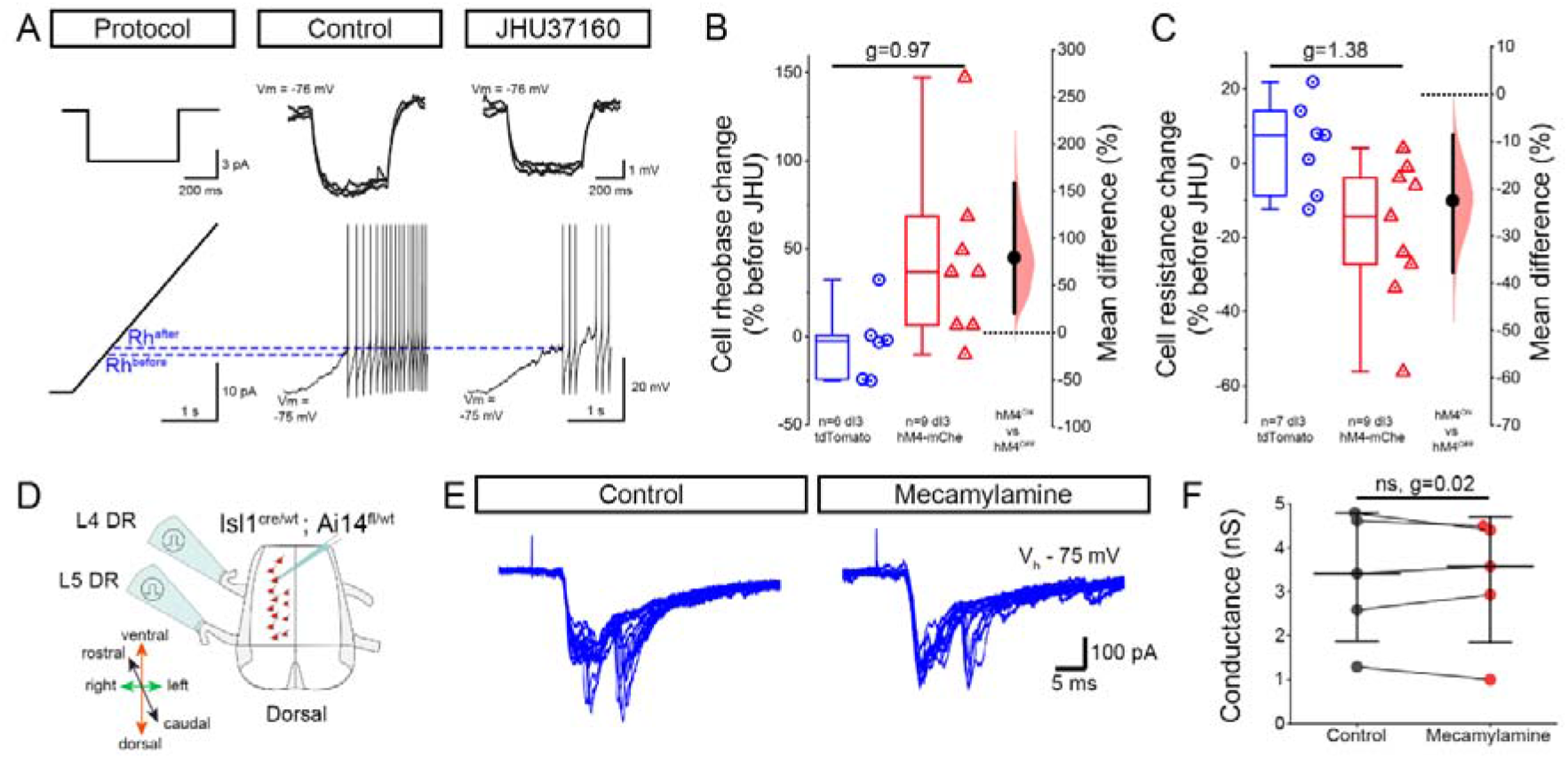
JHU37160 induces a reduction of excitability specifically in dI3 hM4D(Gi)ON while mecamylamine shows no modulation of dorsal root evoked excitatory inputs to dI3 neurons. (A) Step or ramp of current responses from dI3 neurons expressing hM4D(Gi) following injection of AAV2/5-hSyn-Flp^ON^-DIO-HA-hM4Di-mCherry in the lumbar spinal cord in control condition or after perfusion of JHU37160 (230nM) to the bath. (B-C) Change in (B) cell rheobase and (C) cell resistance in dI3 neurons with (dI3 hM4-mChe) or without (dI3 tdTomato) expression of hM4D(Gi) following perfusion of JHU37160 to the bath. (D) Schematic of the spinal cord preparation used to record dorsal root evoked excitation of dI3 neurons. (E) Representative traces of dorsal root evoked excitation in a dI3 neuron in control condition and following addition of mecamylamine to the bath. (F) Amplitude of the dorsal root evoked excitatory responses in control conditions and following addition of mecamylamine (100-200µM). Total of 5 root responses in n=4 dI3 neurons from 4 preparation from N=3 animals. Non-hierarchically bootstrapped paired samples Wilcoxon tests and paired Hedge’s G.

**Supplemental Figure 6:**
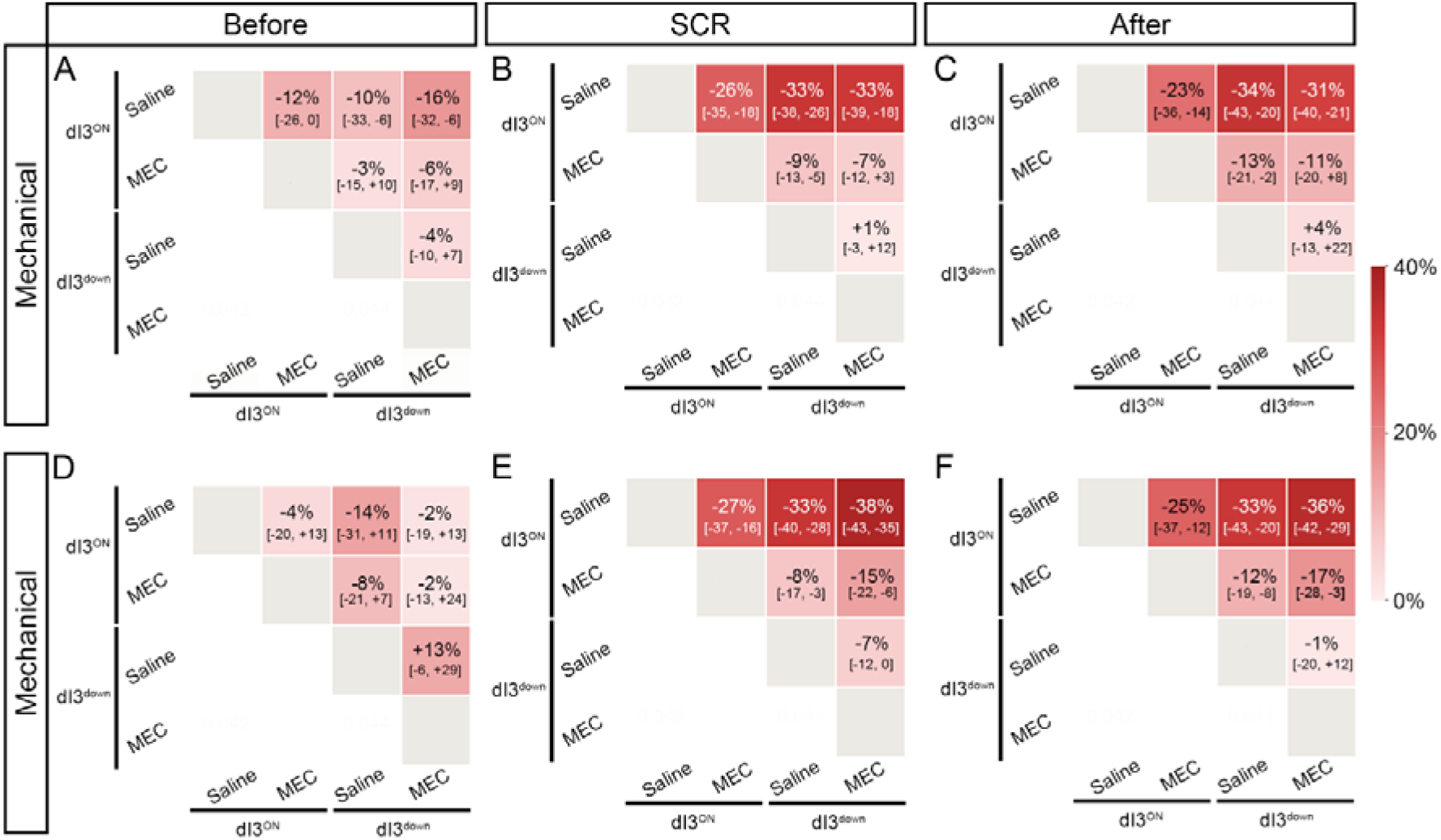
dI3 neurons modulate stumbling correction combining sensory feedback and nicotinic transmission, suggesting a role of Renshaw cell inputs. (A-F) Matrix of the Hodges-Lehmann estimates of the within-animal log ratio back-transformed as percentage changes together with their 95% confidence intervals. Each cell gives the estimate of the column condition relative to the row with colour shading indicating the magnitude of the change. (A-C) mechanical and (D-F) electrical perturbations in the step before, during and after SCR. (A-F) N=7 animals.

***Supplemental video 1:*** Video of the uneven elevated horizontal ladder task for each group tested dI3^ON^ + saline, dI3^down^ + saline, dI3^ON^ + mecamylamine and dI3^down^ + mecamylamine.

***Supplemental video 2:*** Video of the horizontal elevated vibrating beam task for each group tested dI3^ON^ + saline, dI3^down^ + saline, dI3^ON^ + mecamylamine and dI3^down^ + mecamylamine.

***Supplemental video 3:*** Video of stumbling corrective reaction following mechanical perturbation for each group tested dI3^ON^ + saline, dI3^down^ + saline, dI3^ON^ + mecamylamine and dI3^down^ + mecamylamine.

***Supplemental video 4:*** Video of stumbling corrective reaction following mechanical perturbation for each group tested dI3^ON^ + saline, dI3^down^ + saline, dI3^ON^ + mecamylamine and dI3^down^ + mecamylamine.

## Notes

### Competing Interest Statement

Robert M. Brownstone is a co-founder of Sania Therapeutics Inc.

### Summary of Updates

In the revised version of this manuscript, we have included data from a new series of experiments in which we use trans-synaptic rabies virus tracing to positively identify Renshaw cells that are pre-synaptic to dI3 neurons. We also added data describing the monosynaptic inputs received by dI3 neurons from motor axons. In the revision process, we made the decision to move away from using two separate models for the behavioural experiments. That is, we have now excluded the data using mouse genetics alone to manipulate dI3 neurons given the effects of this manipulation on more than the lumbar dI3 neurons. Although the results from the excluded experiments were the same, the specificity of the manipulation was lower, leaving other interpretations open. We therefore conducted an additional series of behavioural experiments in which the expression of DREADD receptors in dI3 neurons was induced via viral manipulation. In this revised version, we have revised the Introduction and extensively rewritten the discussion to clarify our use of control theory terms.

