## Supplementary figures and images for "Intrinsic spinal cord circuits compare sensory inputs and efference copies to correct for perturbations to ongoing movement"

### Supplemental Figure 1

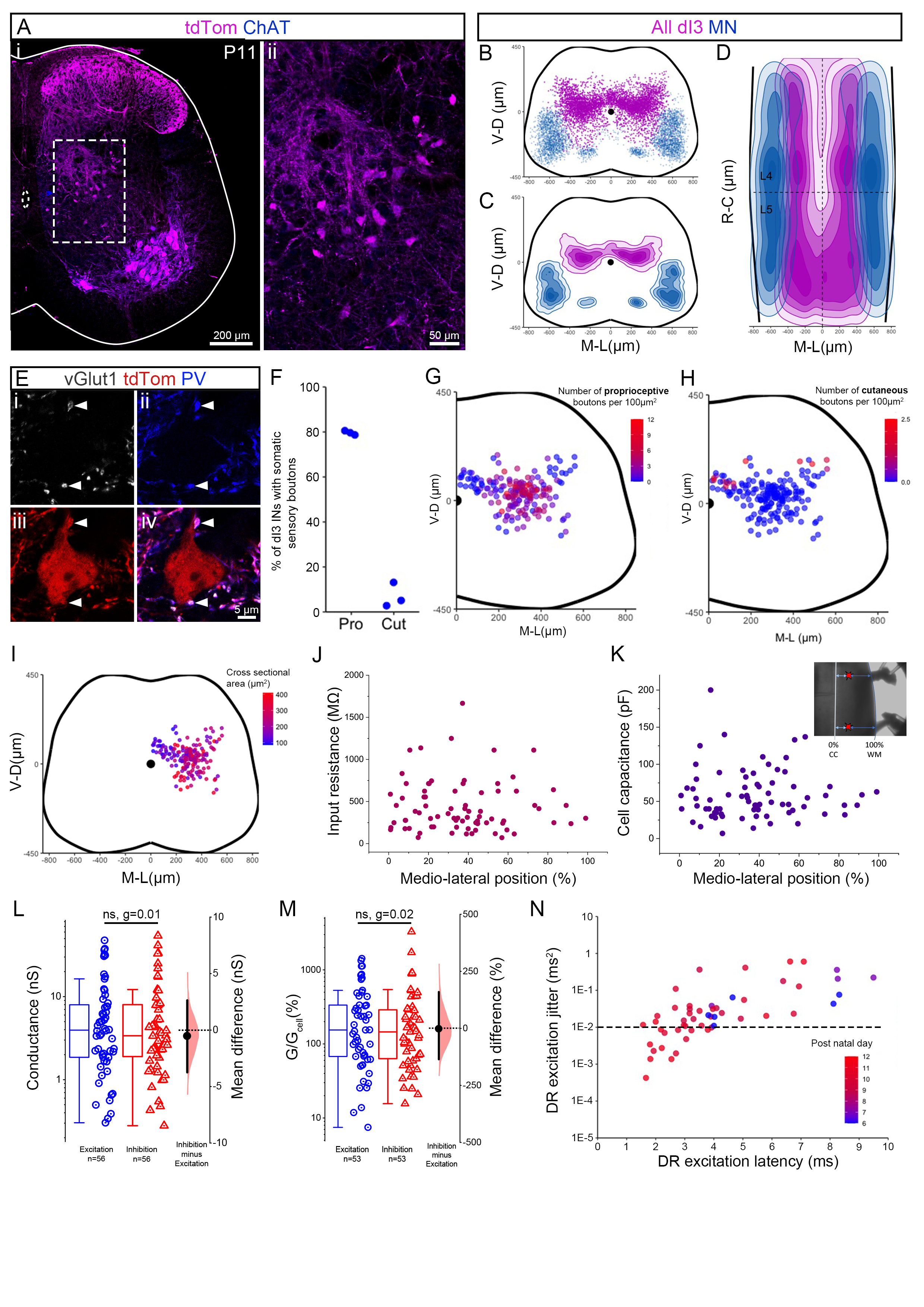

### Supplemental Figure 2

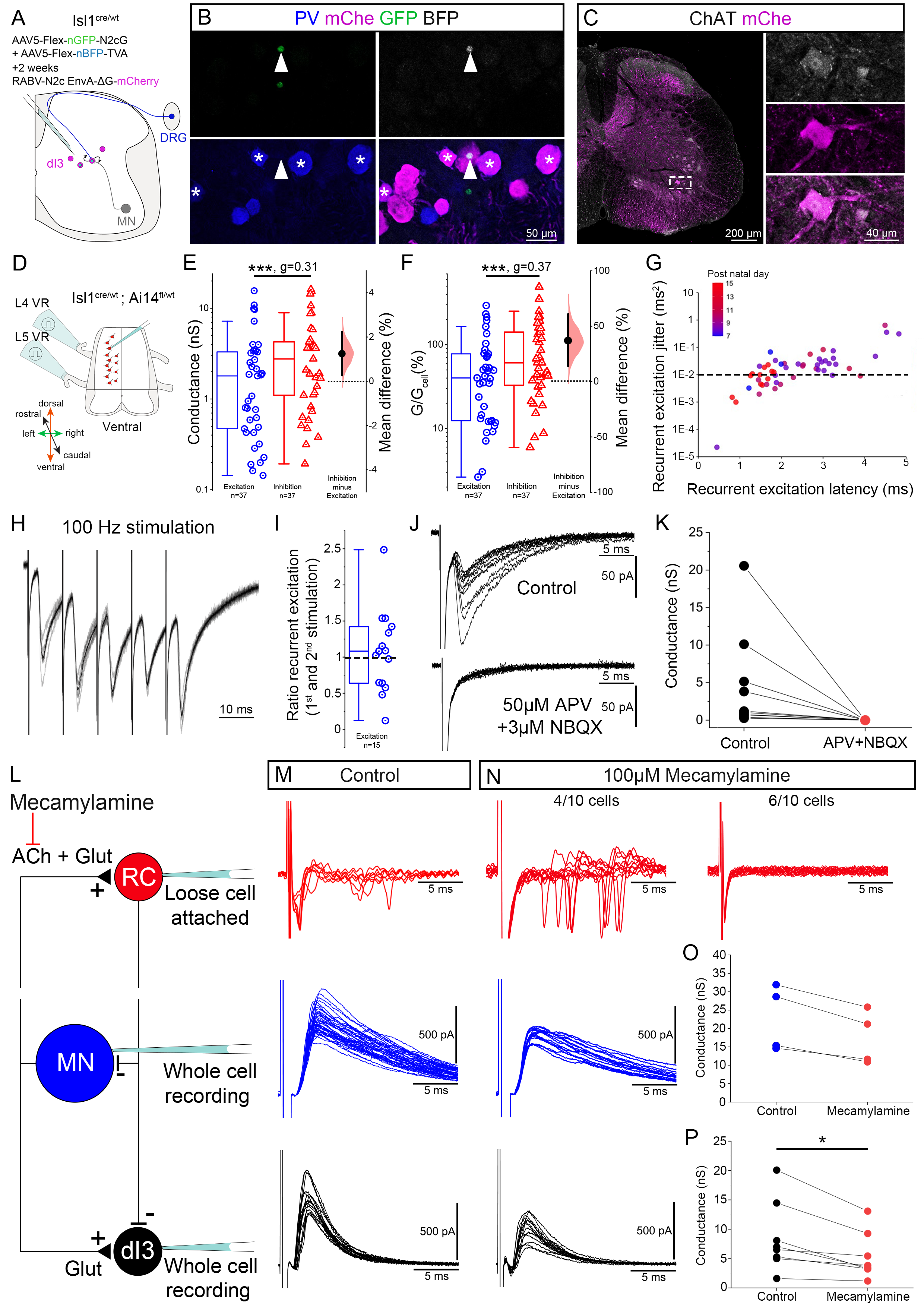

### Supplemental Figure 3

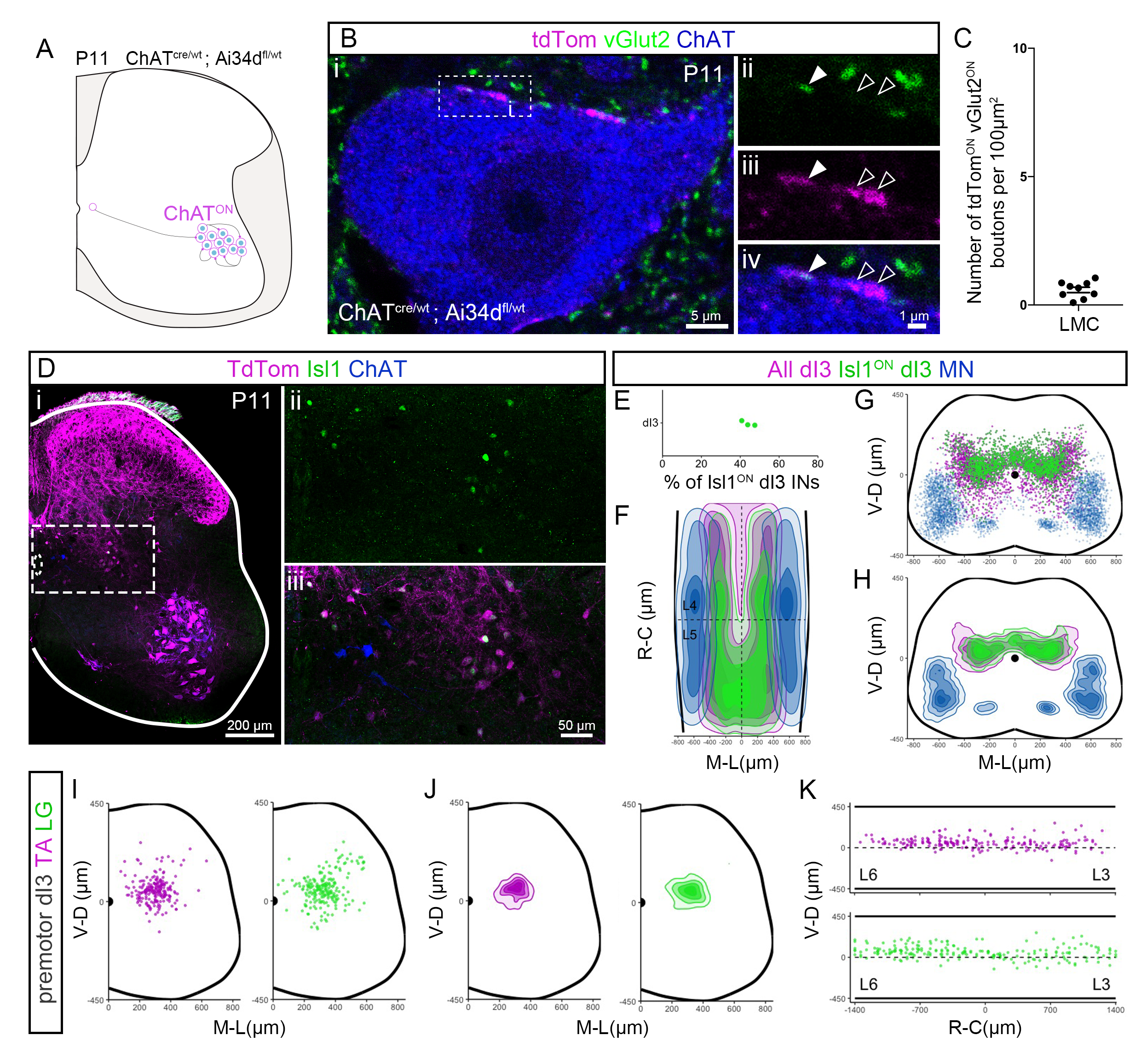

### Supplemental Figure 4

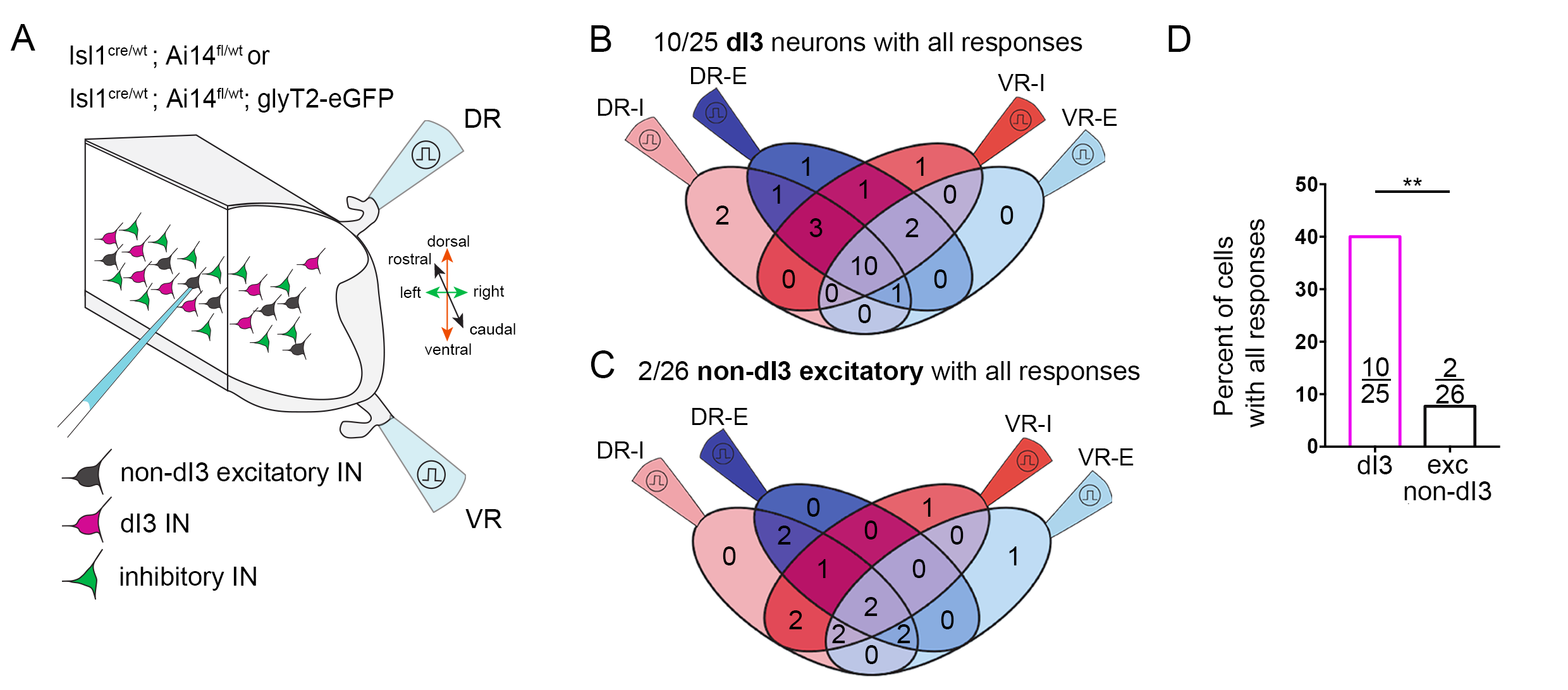

### Supplemental Figure 5

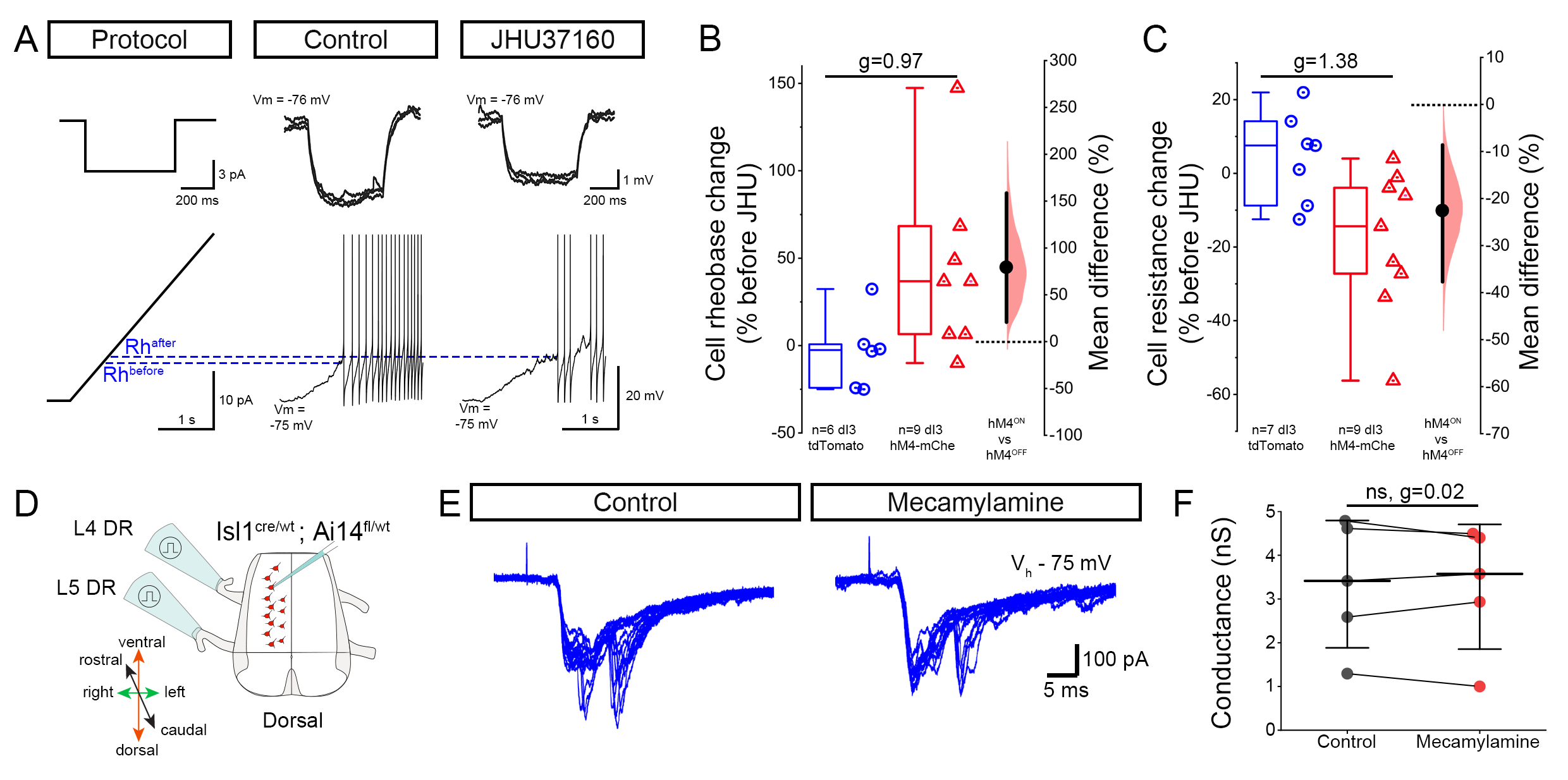

### Supplemental Figure 6

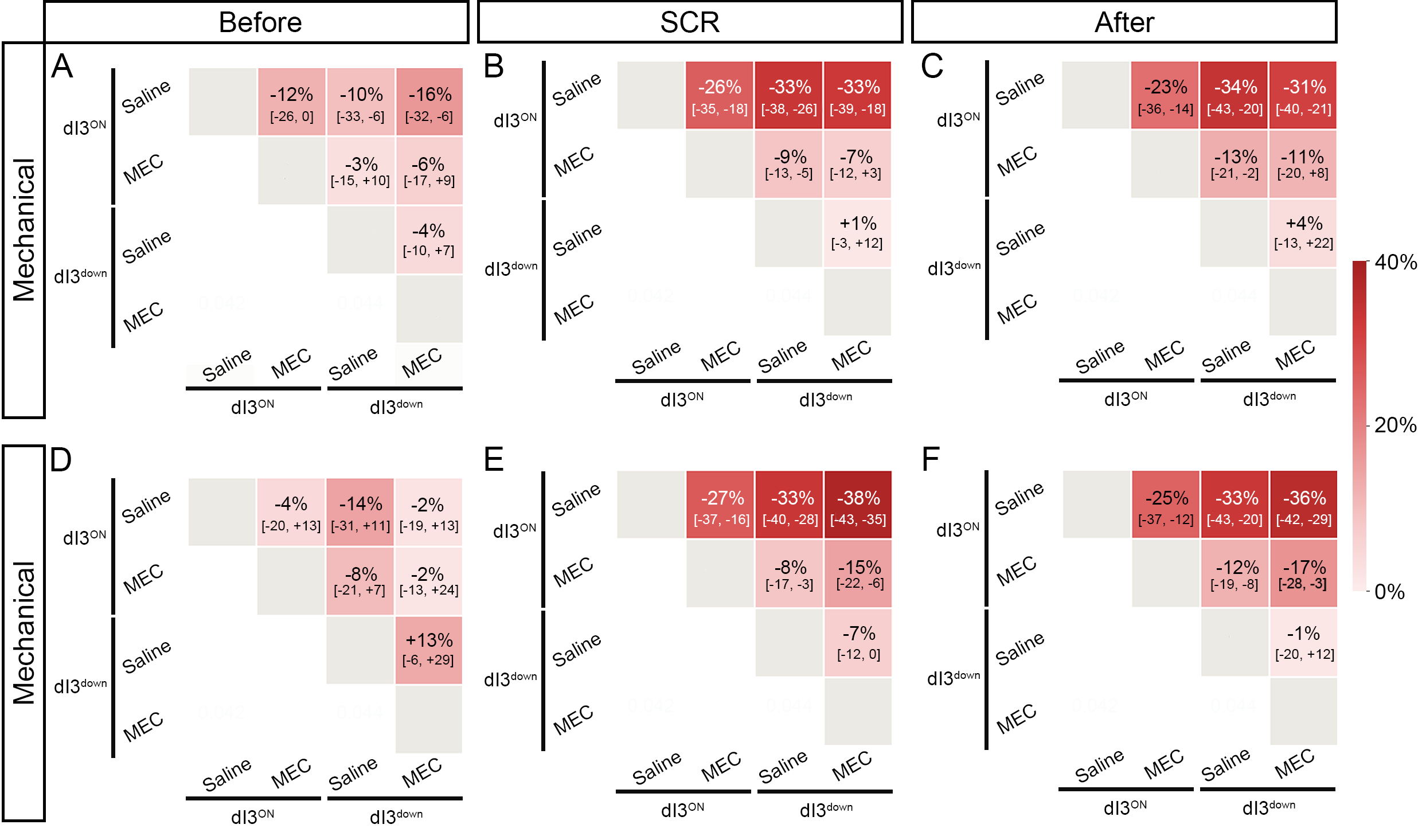
